# Somatic deficiency causes reproductive parasitism in a fungus

**DOI:** 10.1101/2020.08.21.260869

**Authors:** Alexey A. Grum-Grzhimaylo, Eric Bastiaans, Joost van den Heuvel, Cristina Berenguer Millanes, Alfons J.M. Debets, Duur K. Aanen

## Abstract

Some multicellular organisms can fuse because mergers potentially provide mutual benefits. However, experimental evolution in the fungus *Neurospora crassa* has demonstrated that free fusion of mycelia favours cheater lineages, but the mechanism and evolutionary dynamics of dishonest exploitation are unknown. Here we show, paradoxically, that all convergently evolved cheater lineages have similar fusion deficiencies. These mutants are unable to initiate fusion but retain access to wild-type mycelia that fuse with them. This asymmetry reduces cheater-mutant contributions to somatic substrate-bound hyphal networks, but increases representation of their nuclei in the aerial reproductive hyphae. Cheaters only benefit when relatively rare and likely impose genetic load reminiscent of germline senescence. We show that the consequences of somatic fusion can be unequally distributed among fusion partners, with the passive non-fusing partner profiting more. We discuss how our findings may relate to the extensive variation in fusion frequency of fungi found in nature.

## Main

Cheating is an almost universal problem in social systems that depend on cooperation ^1^. For example, in multicellular organisms some cells altruistically support reproduction by genetically related cells ^2–6^. Such reproductive division of labour can be exploited by cheating mutant cells that manage to become overrepresented among the reproductive propagules even when higher-level organismal fitness declines. Experimental evolution approaches have shown that cheating mutants can emerge in simple multicellular organisms ^7–9^, but major open questions on how genetic mechanisms mediate or inhibit the spread of cheating have remained unanswered. Understanding the ubiquity of selection pressure for selfish traits and the social processes that regulate cheating is of paramount importance to appreciate the evolutionary stability of social systems throughout the levels at which social interactions have evolved. Considerable research has focused on cooperation and conflict in prokaryotes and animal societies, but studies in modular multicellular eukaryotes have hardly been done. In this paper, we explore the genetic basis and the operational mechanisms of cheating in a filamentous fungus where mycelia often fuse and cell compartmentalization is limited so that nuclei can migrate across the hyphae.

Filamentous fungi form hyphae that branch and fuse regularly to form a dense, radially growing mycelial network ^10^. Hyphal fusion is a costly somatic trait that pays off because it makes asexual spore production more efficient ^11^. Such spores are produced by aerial hyphae, which can thus be considered as reproductive structures supported by a substrate-bound somatic hyphal network that acquires the organic resources for growth. The cells in most fungal mycelia are incompletely compartmentalized so that nuclei can move freely through most if not all parts of the fungal colony’s mycelial network, including the germline aerial hyphae ^12,13^. Therefore, the cooperating units in a fungal colony are the nuclei, and not the cells as in most other multicellular organisms ^12^. Filamentous fungi can also fuse with other colonies, potentially leading to the formation of chimeras. Cooperative nuclei in such chimeras may then provide fitness benefits to genetically unrelated nuclei, but face the risk that those could be non-cooperative, selected to exploit their partner nuclei for selfish reproductive benefits. Such cheaters are known to exist. Experimental evolution of a *Neurospora crassa* wild-type strain that freely fuses resulted in the emergence of cheaters coexisting with cooperative variants, but a fusion-deficient mutant appeared to preclude the evolution of cheaters ^8^. These cheaters were social parasites ^14^ of the wild type, having an increased probability to become spores relative to the ancestor. They also decreased total spore production both in monoculture and in mixed culture with the competitor, consistent with the general expectations outlined above.

To find putative genes underlying cheating in *N. crassa*, we sequenced the genomes of all identified evolved cheaters and cooperative variants that emerged in our lab cultures, and compared them with their ancestors (Supplementary Tables 1, 2). We found striking parallel evolution among cheaters. In six of the eight independent free-fusion lines the cheater had acquired unique mutations in the gene *soft* (*so*, NCU02794; ^15^), of which five lead to loss of function based on *in silico* predictions (a premature stop-codon, intron-junction mutation, and three frameshift mutations). The cheater of the sixth line had acquired a mutation in *so* leading to an amino acid substitution (L825P). The gene *so* encodes a cytoplasmic protein with unknown molecular function, but has been shown to be involved in cell-cell communication between fusing spore germlings ^16^. *so*-knockout mutants display strongly reduced hyphal and germling fusion compared to the wild type, have reduced spore production, and are female sterile ^17^. Cheaters of the other two lines, which did not have mutations in *so*, had mutations in other known fusion genes – *ham-5* and *ham-8* (NCU01789 and NCU02811, respectively). Knockout mutants of those genes showed similar phenotypic effects on fusion as *so* knockouts ^18,19^. While *ham-8* had acquired a loss-of-function mutation W277* (premature stop-codon formation), *ham-5* had an intronic change, which would not be identified as an inactivating mutation based on *in silico* predictions. However, we confirmed that the *ham-5* cheater has the same phenotype as a clean *ham-5* deletion strain, suggesting that this seemingly neutral intronic mutation also induces a loss of function. The probability that the observed mutations affected fusion genes in all eight independently evolved cheaters by chance is exceedingly low (exact binomial test, *P* = 3.991e-06, see Supplementary Methods). The observed parallelism in loss-of-function mutations in fusion genes therefore indicates that losing the ability to initiate fusion underlies cheating in *N. crassa*.

We used an isogenic deletion mutant of *so* (Δ*so*, Supplementary Table 3) to test this hypothesis, because cheaters also carried additional mutations. We first tested whether disruption of *so* leads to the cheating phenotype, by performing pair-wise competition experiments between the wild type and Δ*so*. By varying the initial frequency of Δ*so* in the competition mixture, we found that Δ*so* indeed has a competitive benefit over the wild type but only at frequencies up to 30% and a disadvantage at higher frequencies (Fig. 1a, Supplementary Figs. 1, 2, see Supplementary Discussion). This result is consistent with the evolution experiment, where cheaters did not go to fixation in seven of the eight lines. Consistent with the characterization as a cheater, total spore production of the mixture decreased when Δ*so* frequency increased (Fig. 1b), corroborating that *so* disruption is causal to the cheating phenotype. To further investigate the link between reduced fusion and cheating we tested seven other known fusion mutants (Δ*mak-1*, Δ*ham-3*, Δ*ham-4*, Δ*ham-5*, Δ*ham-6*, Δ*ham-7*, Δ*ham-8*; Supplementary Figs. 3, 4). Four of these displayed similar cheating behaviour as Δ*so*, i.e. had a competitive benefit at low frequency and a negative effect on spore yield in monoculture (Supplementary Figs. 5, 6). These results further supported a link between reduced fusion and cheating.

**Fig. 1:**
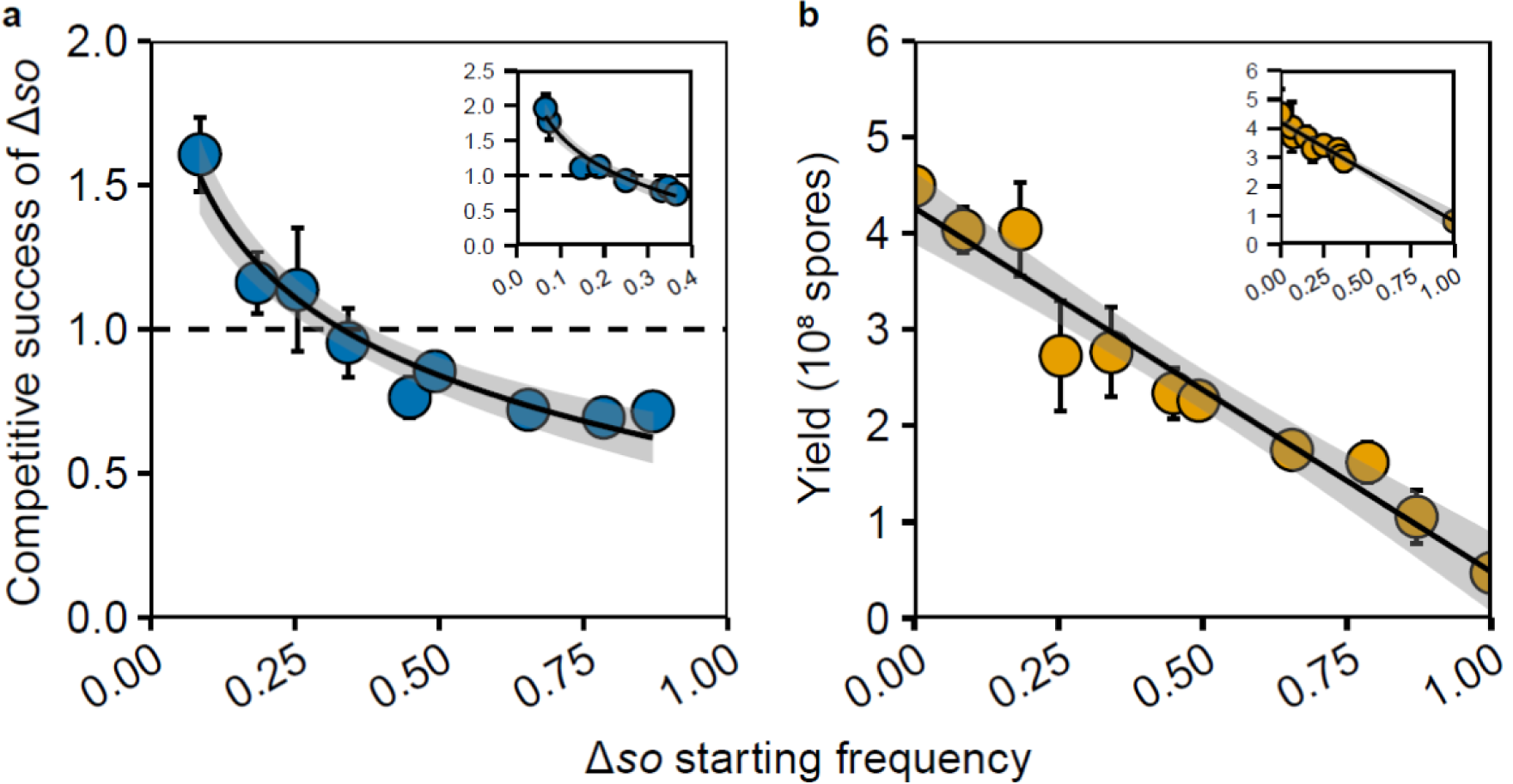
Negative frequency dependence of the benefit of Δ*so* in competition with wild type and the negative effect of Δ*so* on sporulation. **a**, Δ*so* has a competitive advantage relative to wild type at frequencies below ∼30%. **b**, As the frequency of Δ*so* spores in the inoculum increases the total spore production decreases. The inserts in **a** and **b** show the results of a separate experiment testing a narrower range of frequencies around 25%. Shaded areas around the trendlines are 95% confidence areas of the fitted curve. Error bars are 95% confidence intervals calculated from three biological replicates.

Since somatic fusion appeared to be a requirement for cheating, it seemed paradoxical that the cheaters had disrupted fusion genes. We therefore hypothesised that it is the wild-type mycelia that fuse with the cheaters, and that the cheaters profit unilaterally. We first set out to test whether wild type could still fuse with the Δ*so* cheater. Earlier studies showed that the Δ*so* mutant is capable of fusion, albeit at very low frequencies ^17^, but it was still unknown if fusion between wild type and Δ*so* could occur in our experimental setup. We therefore ran transfer experiments with mixtures of Δ*so* and wild type, and measured the fraction of heterokaryotic spores (i.e. containing both Δ*so* and wild-type nuclei) formed by chimeric individuals originating from fusion between wild type and Δ*so*. Our results show that wild type and Δ*so* frequently fuse (Fig. 2a). Already after the first transfer, more than 20% of the spores in Δ*so*/wt mixtures were heterokaryotic, increasing to up to 40% by the fourth transfer. A control experiment where we mixed differently marked Δ*so* revealed almost no heterokaryons (<0.2%). These results demonstrate that Δ*so* cheaters fuse at negligible frequencies with themselves, but wild types frequently fuse with Δ*so* cheaters when Δ*so* is rare, highlighting extreme asymmetry in the fusion trait with the wild type almost always being responsible for fusion with the Δ*so* cheater.

**Fig. 2:**
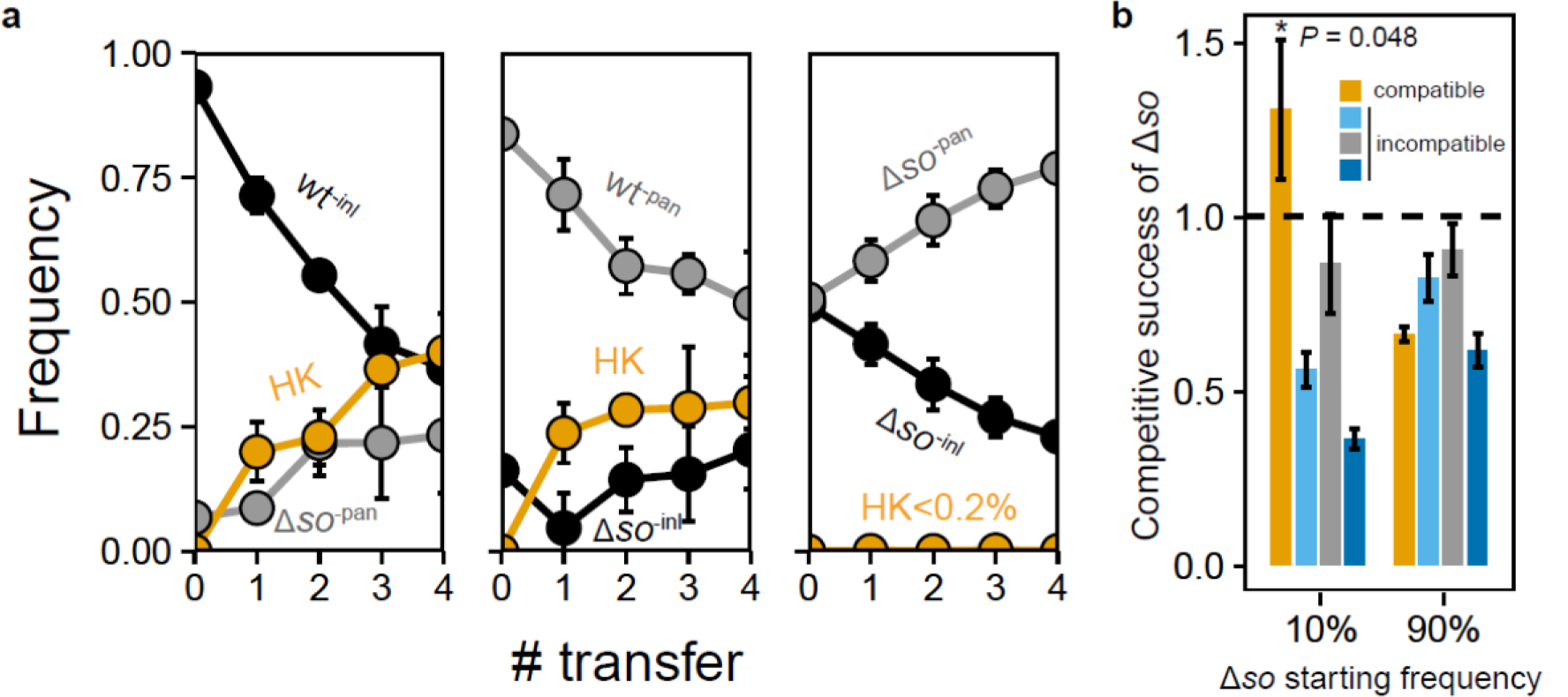
The frequency of fusion between wild type and Δ*so* mycelia and the necessity of successful fusion for Δ*so* to exploit the wild type. **a**, Fusion readily occured between wild type and Δ*so*, but hardly between Δ*so* strains. The first two panels show the results of experiments with reciprocally labelled Δ*so* and wild-type strains, at low Δ*so* frequency (5 to 15%), while a third panel depicts a control experiment with separately labelled Δ*so* mixed in 1:1 initial ratio. *inl*-deficient and *pan*-deficient strains are indicated in black and grey, respectively. The orange line shows the frequency of heterokaryons (HK). **b**, Competition experiments between Δ*so* and compatible (orange bars) and three nearly-isogenic incompatible wild-type strains at two starting frequencies (10% and 90%). Δ*so* only increased when grown with a compatible competitor and only at low frequencies (one-sided *t*-test, t-value = 2.986799, df = 2, *P* = 0.0480968). Error bars are 95% confidence intervals based on three biological replicates.

Having demonstrated frequent fusion between passive Δ*so* cheaters and active wild types, we then tested whether fusion is required for cheating. To exclude the possibility of fusion we used near-isogenic somatically incompatible wild types as competitors against the Δ*so* cheater. Somatic incompatibility prevents chimera formation by inducing cell death if the nuclei in a fused cytoplasm carry different alleles at somatic-incompatibility loci ^20^. This showed that Δ*so* cheaters did not benefit in competition with incompatible wild types, neither at low nor at high frequencies (Fig. 2b, Supplementary Fig. 7). Successful fusion is thus necessary for Δ*so* to become an exploiter of the wild type.

We then asked how the Δ*so* cheater can realize a relative benefit within the chimeric mycelium. We hypothesised that hyphae with a high proportion of wild-type nuclei will be more inclined to fuse and thus have a higher tendency to perform supportive somatic functions ^11,13^. In contrast, hyphae with more Δ*so* nuclei should have reduced probability to fuse, and will thus have a higher probability to be part of aerial reproductive hyphae that become spores ^21^. This hypothesis predicts that the frequency of Δ*so* cheater nuclei should not increase during mycelial growth and that Δ*so* nuclei should increase their representation in the spores compared to the parental mycelium. qPCR analyses of ten randomly selected chimeric individuals with variable frequencies of Δ*so* showed that the frequency of Δ*so* had decreased after colony growth (Fig. 3a, one-sided *t-*test, t-value = −11.692, df = 9, *P* = 4.803e-07; Supplementary Fig. 8, see Supplementary Discussion), irrespectively of the starting frequency (linear regression, F-statistic 1,8 = 5.309, df = 8, *P* = 0.05014), indicating that the fusion-deficient mutant does not bias its representation during somatic growth of a chimera. However, we found that Δ*so* nuclei become overrepresented in the spores, provided the frequency of Δ*so* in the chimeric mycelium remained below 60%. Conversely, Δ*so* nuclei were underrepresented in the spores when they reached high frequencies in the chimeras (Fig. 3b, linear regression, *P* < 0.001; Supplementary Fig. 8, see Supplementary Discussion). These combined results confirmed that active hyphal fusion is indeed linked with supportive somatic functions and that fusion-deficient mutants exploit this asymmetry to obtain a relative advantage in the germline. This benefit was negatively frequency dependent, consistent with the finding that cheaters and wild types stably coexisted in all but one of the evolution lines of *N. crassa* ^8^.

**Fig. 3:**
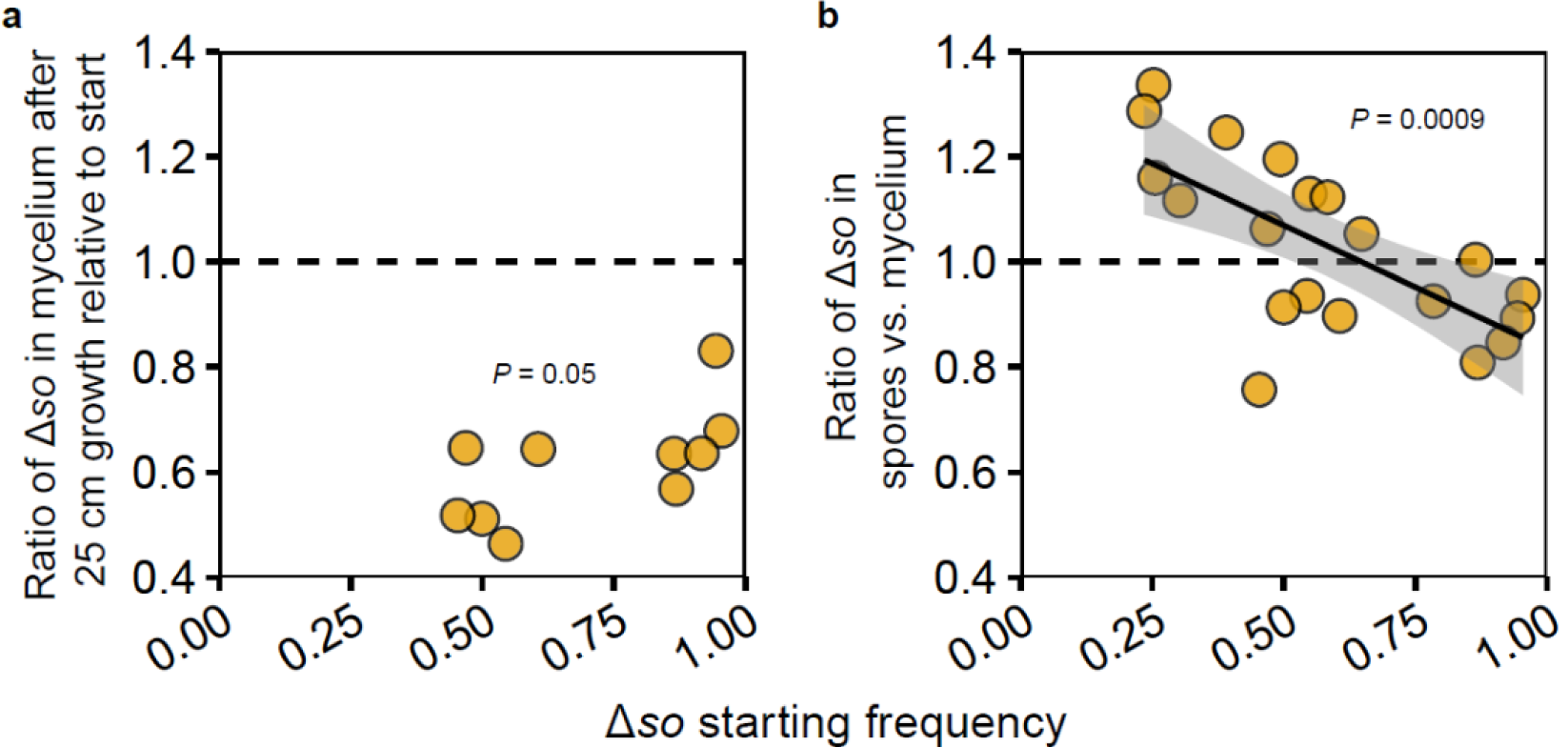
How Δ*so* realizes a competitive benefit in chimeras. **a**, During mycelial growth, the frequency of Δ*so* decreased (one-sided *t-*test, t-value = −11.692, df = 9, *P* = 4.803e-07), irrespectively of the starting frequency (linear regression, F-statistic 1,8 = 5.309, df = 8, *P* = 0.05014). **b**, In contrast, during spore formation Δ*so* nuclei have a benefit over wild-type nuclei as long as starting frequencies remain below ∼60%. The shaded area around the trendline is the 95% confidence area.

Figure 4 presents a synthesis of our results, illustrating why cheating is negatively frequency dependent, and resolves the paradox that a fusion-deficient mutant has a relative benefit over a freely-fusing competitor in spite of fusion being required to reap this benefit. While it has been shown before that the costs and benefits of fusion can be asymmetrically distributed between fusing partners ^8,22,23^, our present results imply that the very ability to fuse is a cause of asymmetrical distribution of these costs and benefits. The fusion-deficient mutants benefit from wild-type strains initiating fusion while never taking such initiative themselves. This demonstrates that hyphal fusion is an altruistic trait, because it facilitates efficient resource distribution and utilization during somatic growth to the benefit of all aerial hyphae that produce and disperse spores during asexual reproduction ^21^. Since cheaters have a lower chance to participate in hyphal fusion, most of fusion occurs between hyphae with many wild-type nuclei. Consequently, less-well connected cheater-rich hyphae have a higher probability to become aerial hyphae and to contribute to spore formation.

**Fig. 4:**
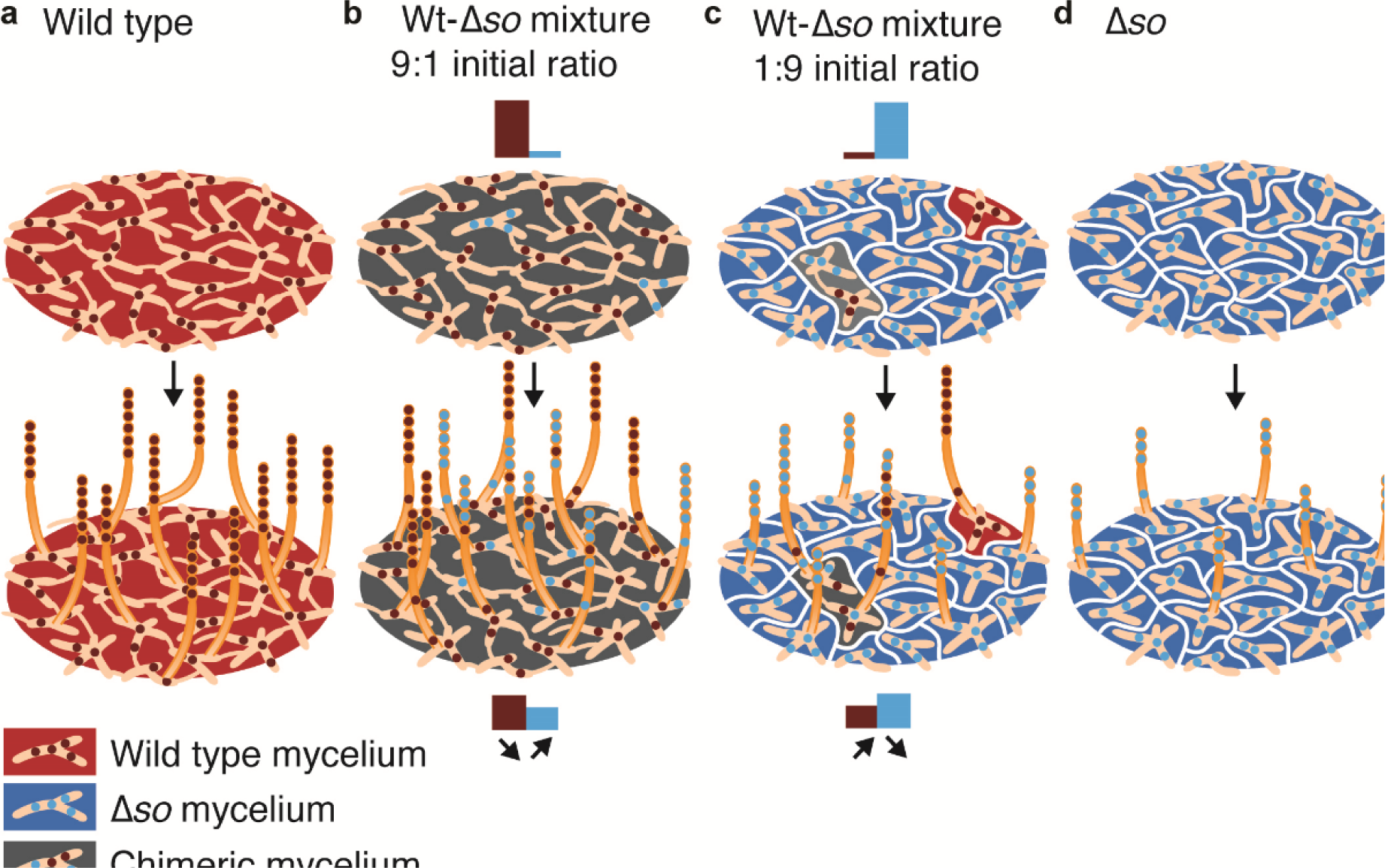
Frequency-dependent selection of the Δ*so* cheater and consequences for spore production. **a**, In a wild-type colony, frequent hyphal fusion allows efficient resource distribution within a mycelial network for optimal development of reproductive aerial hyphae. **b**, At low frequency, Δ*so* cheaters are surrounded by connected wild-type hyphae, and have a high chance that at least one wild-type hypha will fuse with them. This gives cheaters access to well-connected wild-type mycelia, and since the cheater nuclei make hyphae less likely to participate in fusion they have a higher likelihood to become overrepresented in the reproductive aerial hyphae. **c**, However, when Δ*so* cheaters reach high frequencies the culture becomes fragmented (white borderlines) and Δ*so* mycelia will often remain unconnected to wild-type hyphae. This reduces total spore yield, and provides a relative benefit to wild-type patches that remain isolated from Δ*so*. **d**, A Δ*so* monoculture sporulates very poorly because the somatic structures are maximally fragmented and unable to efficiently mobilize resources to support the reproductive aerial hyphae. The bars indicate the relative amounts of spores of Δ*so* cheater and wild type.

The convergent emergence of cheaters with the same or similar genetic deficiencies in our evolution experiments shows that they arise often and have sufficiently high selection coefficients to proliferate. Our results imply that the morphological mutants frequently found in lab strains of *N. crassa* are most likely products of unintended selection of *so*-cheater variants during routine propagation ^24^. Maintaining live cultures by serial transfers, instead of using frozen stocks, may thus reduce overall performance, which begs the question how common or rare cheating mutants are in nature. The high inoculation density of clonally related spores that we used in our evolution experiments may resemble the natural growth conditions of *N. crassa* because it preferentially grows in post-fire habitats ^25^. The sexual spores are heat tolerant and germinate upon heat activation. The emerging mycelium then rapidly colonizes empty niche space likely via massive local dispersal of asexual spores. If fusion-deficient mutants accumulate during this natural asexual proliferation, it would be reminiscent to senescence of an aging multicellular individual. Fusion mutants may survive fire events if they end up in homokaryotic sexual spores. However, the germinating spores of fusion-deficient cheaters will remain dependent on fusion with wild-type colonies, which will likely be somatically incompatible because of segregated incompatibility alleles in sexual offspring ^20^. Genetic load imposed by fusion-deficient mutants may therefore rarely persist across seasons in wild *N. crassa* populations.

Specialization on post-fire habitats constrains the asexual life span of *N. crassa* in contrast to many fungi that lack environmentally-restricted life spans. Indeed, some fungal colonies are among the longest-lived organisms as for example *Armillaria* species with an estimated age of millennia ^26^. Such extreme longevity requires efficient policing mechanisms against cheating mutants, which may be associated with reduced mutation rates ^27,28^. Additionally, many fungi have stronger cell compartmentalization than *N. crassa* and synchronized nuclear division, mechanisms that both restrict dispersal of selfish nuclei through mycelia ^12,29^. Restricting fusion via allorecognition has also been recognized as a mechanism to reduce the risk of parasitism ^30–32^. Reducing fusion frequency itself is yet another way to prevent cheating, and the extensive variation among fungi in fusion frequency, reflected in their tendency to form physically separated mutant sectors within a clonal mycelial colony ^33^, may thus generally reflect different sensitivities to cheating mutants depending on species-specific ecology. Our experiments demonstrate that fusion is best conceptualized as an altruistic trait that can be exploited to a degree set by the environmental conditions such as extrinsic mortality, which in turn will likely select for defence mechanisms.

Buss ^12^ argued that animals and fungi are more sensitive to exploitation than plants, since plants have rigid cell walls limiting their mobility, while animal cells and fungal nuclei can disperse within individuals. However, in fungi the opportunities for dispersal also depend on the degree of modular connectivity ^13^, which does not apply to unitary animals. It seems clear that development of a multicellular organism by non-clonal cell aggregation should give more scope for parasitism than clonal development from a single-celled zygote ^1,2,7,34^. However, individuals with clonal development are not immune to social parasitism since nuclei can mutate and be transmitted to other individuals via fusion. Modular organisms such as fungi that lack early germline sequestration are particularly sensitive, since all body parts can ultimately reproduce. However, even early germline sequestration does not guarantee immunity and still leaves the potential for social parasitism via fusion. A striking example from humans came to light decades ago ^35^ when genetic analysis showed that a woman could not be the biological parent of any of her four presumed children, while her parents were confirmed to be the grand parents of these children. The only reasonable explanation for this enigma was that the mother was a chimera, i.e. the product of a fusion of two full-sister embryos, of which one had contributed the gonads and the other the somatic cells.

## Methods

please see Supplementary Information.

## Data availability

Raw sequencing data was deposited at NCBI BioProject number PRJNA605104. The web-link for reviewers: https://dataview.ncbi.nlm.nih.gov/object/PRJNA605104?reviewer=s2h97d05ubpjqdemfgv2d4o20l.

## Code availability

Computer code used in the analysis is available on request from the corresponding author.

## Acknowledgements

We thank Jacobus Boomsma, Ben Auxier, Bas Zwaan for helpful comments on this manuscript, and Marc Maas for making Figure 4. DKA, EB and AAG-G were supported by the Netherlands Organisation for Scientific Research (VICI; NWO 86514007).

## Authors contributions

DKA, AAG-G, AJMD, EB developed the concept. AAG-G, EB, CBM performed the experiments. JvdH developed the bioinformatics pipeline. AAG-G analysed the genomes. DKA and AAG-G wrote the manuscript with contributions from EB and AJMD.

## References

1. Velicer, G. J., Kroos, L. & Lenski, R. E. Developmental cheating in the social bacterium Myxococcus xanthus. Nature 404, 598–601 (2000).

2. Fisher, R. M., Cornwallis, C. K. & West, S. A. Group formation, relatedness, and the evolution of multicellularity. Curr. Biol. 23, 1120–1125 (2013).

3. West, S. A. & Cooper, G. A. Division of labour in microorganisms: an evolutionary perspective. Nat. Rev. Microbiol. 14, 716–723 (2016).

4. Ratcliff, W. C., Denison, R. F., Borrello, M. & Travisano, M. Experimental evolution of multicellularity. Proc. Natl. Acad. Sci. U. S. A. 109, 1595–1600 (2012).

5. Ratcliff, W. C., Fankhauser, J. D., Rogers, D. W., Greig, D. & Travisano, M. Origins of multicellular evolvability in snowflake yeast. Nat. Commun. 6, 6102 (2015).

6. Grosberg, R. K. & Strathmann, R. R. The evolution of multicellularity: a minor major transition? Annu. Rev. Ecol. Evol. Syst. 38, 621–654 (2007).

7. Kuzdzal-Fick, J. J., Fox, S. A., Strassmann, J. E. & Queller, D. C. High relatedness is necessary and sufficient to maintain multicellularity in Dictyostelium. Science (80-.). 334, 1548–1551 (2011).

8. Bastiaans, E., Debets, A. J. M. & Aanen, D. K. Experimental evolution reveals that high relatedness protects multicellular cooperation from cheaters. Nat. Commun. 7, 11435 (2016).

9. Velicer, G. J., Kroos, L. & Lenski, R. E. Loss of social behaviors by Myxococcus xanthus during evolution in an unstructured habitat. Proc. Natl. Acad. Sci. U. S. A. 95, 12376–12380 (1998).

10. Glass, N. L., Rasmussen, C., Roca, M. G. & Read, N. D. Hyphal homing, fusion and mycelial interconnectedness. Trends Microbiol. 12, 135–141 (2004).

11. Bastiaans, E., Debets, A. J. M. & Aanen, D. K. Experimental demonstration of the benefits of somatic fusion and the consequences for allorecognition. Evolution (N. Y). 69, 1091–1099 (2015).

12. Buss, L. W. The evolution of individuality. (Princeton Univ. Press, Princeton, NJ, 1987).

13. Roper, M., Simonin, A., Hickey, P. C., Leeder, A. & Glass, N. L. Nuclear dynamics in a fungal chimera. Proc. Natl. Acad. Sci. U. S. A. 110, 12875–12880 (2013).

14. Strassmann, J. E. & Queller, D. C. Evolution of cooperation and control of cheating in a social microbe. Proc. Natl. Acad. Sci. U. S. A. 108, 10855–10862 (2011).

15. Wilson, J. F. & Dempsey, J. A. A hyphal fusion mutant in Neurospora crassa. Fungal Genet. Rep. 46, 25–30 (1999).

16. Fleissner, A., Leeder, A. C., Roca, M. G., Read, N. D. & Glass, N. L. Oscillatory recruitment of signaling proteins to cell tips promotes coordinated behavior during cell fusion. Proc. Natl. Acad. Sci. U. S. A. 106, 19387–19392 (2009).

17. Fleißner, A. et al. The so locus is required for vegetative cell fusion and postfertilization events in Neurospora crassa. Eukaryot. Cell 4, 920–930 (2005).

18. Dettmann, A., Heilig, Y., Valerius, O., Ludwig, S. & Seiler, S. Fungal communication requires the MAK-2 pathway elements STE-20 and RAS-2, the NRC-1 adapter STE-50 and the MAP kinase scaffold HAM-5. PLoS Genet. 10, e1004762 (2014).

19. Fu, C., Ao, J., Dettmann, A., Seiler, S. & Free, S. J. Characterization of the Neurospora crassa cell fusion poteins, HAM-6, HAM-7, HAM-8, HAM-9, HAM-10, AMPH-1 and WHI-2. PLoS One 9, e107773 (2014).

20. Aanen, D. K., Debets, A. J. M., Glass, N. L. & Saupe, S. J. Biology and genetics of vegetative incompatibility in fungi. in Cellular and molecular biology of filamentous fungi 274–288 (American Society of Microbiology, 2010).

21. Springer, M. L. Genetic control of fungal differentiation: the three sporulation pathways of Neurospora crassa. BioEssays 15, 365–374 (1993).

22. Pittenger, T. H. & Brawner, T. G. Genetic control of nuclear selection in Neurospora heterokaryons. Genetics 46, 1645–1663 (1961).

23. Vreeburg, S., Nygren, K. & Aanen, D. K. Unholy marriages and eternal triangles: how competition in the mushroom life cycle can lead to genomic conflict. Philos. Trans. R. Soc. B Biol. Sci. 371, 20150533 (2016).

24. Kafer, E. Improved backcrossed strains giving consistent map distances. Fungal Genet. Rep. 29, 41–44 (1982).

25. Jacobson, D. J. et al. New findings of Neurospora in Europe and comparisons of diversity in temperate climates on continental scales. Mycologia 98, 550–559 (2006).

26. Smith, M. L., Bruhn, J. N. & Anderson, J. B. The fungus Armillaria bulbosa is among the largest and oldest living organisms. Nature 356, 428–431 (1992).

27. Anderson, J. B. et al. Clonal evolution and genome stability in a 2500-year-old fungal individual. Proc. R. Soc. B Biol. Sci. 285, 1–6 (2018).

28. Hiltunen, M., Grudzinska-Sterno, M., Wallerman, O., Ryberg, M. & Johannesson, H. Maintenance of high genome integrity over vegetative growth in the fairy-ring mushroom Marasmius oreades. Curr. Biol. 29, 2758–2765.e6 (2019).

29. Roberts, S. E. & Gladfelter, A. S. Nuclear autonomy in multinucleate fungi. Curr. Opin. Microbiol. 28, 60–65 (2015).

30. Debets, A. J. M. & Griffiths, A. J. F. Polymorphism of het-genes prevents resource plundering in Neurospora crassa. Mycol. Res. 102, 1343–1349 (1998).

31. Heller, J. et al. Characterization of greenbeard genes involved in long-distance kind discrimination in a microbial eukaryote. PLoS Biol. 14, e1002431 (2016).

32. Gonçalves, A. P. et al. Allorecognition upon fungal cell-cell contact determines social cooperation and impacts the acquisition of multicellularity. Curr. Biol. 29, 3006–3017.e3 (2019).

33. Pontecorvo, G. & Gemmell, A. R. Colonies of Penicillium notatum and other moulds as models for the study of population genetics. Nature 154, 532–534 (1944).

34. Maynard Smith, J. & Szathmáry, E. The major transitions in evolution. (Freeman, Oxford, 1995).

35. Mayr, W. R., Pausch, V. & Schnedl, W. Human chimaera detectable only by investigation of her progeny. Nature 277, 210–211 (1979).

