## Supplementary Information for "Somatic deficiency causes reproductive parasitism in a fungus"

**Supplementary Information to** **“*Somatic deficiency causes reproductive parasitism in a fungus*”**

1. **Supplementary Methods**
2. **Supplementary Discussion**
3. **Supplementary Figures 1-8**
4. **Supplementary Tables 1-3**
5. **Supplementary References 36-50 (continued from the main text)**
1. **Supplementary Methods**

**Strains and routine sub-culturing:** The wild-type *Neurospora crassa* strains as well as single-gene knockout mutants (from the *Neurospora* Functional Genomics Project,^36^) were obtained from the Fungal Genetics Stock Center (RRID:SCR_008143), Manhattan, Kansas, USA (FGSC,^37^). Additional strains carrying *inl* or *pan-2* (or “*pan”* for short) auxotrophic markers (deficiencies for inositol and pantothenic acid, respectively) in the background of the standard laboratory strain were obtained from ^8^. The morphotypes that arose after the evolution experiment in ^8^ were used for the whole genome sequencing (see section “Genomes sequencing and analyses”). Supplementary Table 3 lists the strains used in the study (except morphotypes from the evolution lines). For all competitions and transfer experiments we used strains carrying the *mat A* idiomorph. The strains were kept as spore stocks either on silica gel at +4°C, or in glycerol/peptone (25%/7%) suspension at -80°C. They were routinely propagated on slants with Vogel’s minimal medium (VMM) solidified with 2% agar, and containing 2% sucrose as a carbon source^38^. Inositol and/or pantothenic acid (final concentration 50 mg/l) were added to the media to propagate the respective auxotrophic mutants. All incubations were performed at 25°C with 12/12 hour light/dark regime.

**Counting plates:** Counting plates (or “sorbose-VMM”) were prepared by adjusting VMM, by substituting sucrose for L(-)sorbose (Calbiochem SDS) with the addition of 0.05% glucose and fructose. This medium keeps the single-spore colonies restricted to a small size (~1 cm after a week growth), making feasible counting colonies (up to ~100) on a 9-cm Petri plate. On this medium after 7 days, wild-type colonies produced bright orange heavily-sporulating colonies, while most fusion mutants studied here (Δ*so*, Δ*ham-5*, Δ*ham-6*, Δ*ham-7*, Δ*ham-8*) produced pale-orange, poorly-sporulating flat colonies. Δ*ham-3*, Δ*ham-4*, produced larger but transparent colonies often with a mycelial rim. Δ*mak-1* fusion mutant formed very small (a few mm) transparent colonies. These phenotypes allowed us to distinguish the mutants from the wild type on counting plates (Supplementary Fig. 4).

**Sexual crosses:** Sexual crosses were performed on slants with synthetic crossing medium (SCM) containing 2% sucrose as a carbon source^38^. As most mutants used in this study were female sterile^39^, we used them as males in sexual crosses as follows. Some asexual spores of the knockout mutants were spread onto the 1 d-old recipient female colony. The fertilized culture was incubated until the development of fruit bodies (for 2–3 weeks). The sexual spores were then shot out of the fruit bodies and became visible on the inside of a glass tube as a black film. The sexual spores were collected with a wet cotton swab and were given the required heat shock for 30 min at 61°C to activate germination, but also to kill the residual asexual spores and mycelial fragments. After the heat shock, the sexual spores were spread onto the counting plates supplemented with hygromycin B (final concentration 200 μg/ml). Since single-gene knockouts were constructed by substituting the gene of interest with a hygromycin cassette, the mutants are resistant to hygromycin, making it a selectable marker^36^. A control plate without hygromycin was used to check whether the re-constituted wild-type strain, which lacks hygromycin resistance, would grow along with the rest of the progeny. The candidate progeny colonies were transferred into slants with VMM, checked for the phenotype, tested for the *inl* or *pan* deficiency, and for the mating type. The mating type was determined by inoculating asexual spores onto 3–5 d-old colony (acted as female) of tester strains (Δ*fl*::*Hyg*^r^; mat A and Δ*fl*::*Hyg*^r^; mat a) and checked for the fruit bodies formation at the inoculum spots. Once the mutant was confirmed and the mating type determined, the glycerol/peptone (25%/7%) spore stock was prepared and stored at -80°C until use.

**DNA extraction:** The morphotypes (i.e. cheaters and social types) of *Neurospora crassa* that emerged after our evolution experiment from ^8^, as well as the ancestors, were used for full-genome sequencing. To obtain a large amount of mycelium, the strains were grown in 100 ml MY medium (in 500 ml Erlenmeyer’s flasks, malt extract – 17 g/l, yeast extract 1 g/l) on a rotary shaker (220 rpm) at 25°C for 1–2 days. A few grams wet weight mycelium was collected and stored frozen at -20°C. Upon genomic DNA (gDNA) extraction, the mycelium was frozen in liquid nitrogen and ground to powder by a sterile pestle in a sterile mortar. The gDNA was extracted using the CTAB-based protocol as in ^40^, with minor modifications (such as reduced mass of starting material to 1 g wet mycelium, and not performing RNase A treatment). The quality and quantity of the gDNA were verified on 0.6% agarose gels stained with ethidium bromide, but also using a Nanodrop 2000 (Thermo Fisher Scientific) and Qubit Fluorometer dsDNA assays (Thermo Fisher Scientific).

For qPCRs analyses, the gDNA from mycelium and spores was isolated with DNeasy Plant Mini Kit (Qiagen) following the manufacturer’s instructions.

**Genome sequencing and analyses:** To find mutations, the genomes of the evolved morphotypes (n=36, at ~25x coverage) and the respective ancestors (n=2, at ~40x coverage) of the evolution experiment were sequenced with the Illumina technology (150 bp paired-end reads, HiSeq4000 platform) at BGI (Hong-Kong). The raw reads were filtered and trimmed using the trim-fastq.pl script (--min-length 70 --quality-threshold 30) of the popoolation v.1.2.2 package^41^, to ensure only high-quality reads were used for the subsequent mapping. The filtered reads were mapped to the reference genome sequence of *Neurospora crassa* OR74A (version NC12 + mitochondrial “Supercontig_10.21” contig -> 41102378 nt total with the predicted 9758 protein-coding genes; *Neurospora crassa* Database, RRID:SCR_001372, FungiDB, RRID:SCR_006013) with bwa -mem version 0.7.17-r1188^42^. The SAM files and quality-filtered (mapping quality >= 20) sorted BAM files were generated with samtools 1.9^43,44^. The alignment statistics and the average coverage of reads were calculated using “samtools flagstat” and “samtools depth” commands (Supplementary Table 1). Pairwise (ancestor/evolved) mpileup files were generated with “samtools mpileup -Bf” command. Mpileup files were used as inputs for VarScan v.2.4.4^45^ to call high-confident SNPs and small indels (<4 bp), using “mpileup2cns --min-coverage 10 --min-var-freq 0.8 --p-value 0.005 --variants --strand-filter 0 --output-vcf 1”. The resulting VCF files were inspected for genetic differences by pairwise comparisons, using vcfR v.1.8.0 R package^46^, for which we used a minimum allele frequency difference of 0.8 and a minimum coverage of 10 to filter SNPs and indels (Supplementary Table 2). This pipeline did not yield variants for a couple of social variants (genomes of the morphotypes 11t1 and 14t1), hence we treat those genomes as heterokaryotic, and did not analyse them with more relaxed parameters. The detected variants were visually confirmed in the IGV genome browser v. 2.3.94^47,48^.

To find larger indels, we used unfiltered BAM files and filtered out all reads with soft and hard clipped reads, as well as indel and deletion calls. Furthermore, we filtered on flags 67, 131, 115, 179, 81, 161, 97, 145, 65, 129, 113 and 177, which potentially indicate large indels or deletions or chromosomal rearrangements. Using a custom R 3.6.1^49^ script, we quantified coverage of all these reads in 100 bp windows. The resulting coverage distribution was divided to the total coverage of the initial unfiltered BAM files, which therefore yielded the frequencies of ‘alternate call’ mapped reads to those with ‘normal’ mapping. These frequencies were then compared between all pairwise samples, ordered upon frequency and visually inspected using IGV without any a priori cut-offs. Half of the variants detected using this custom-script method showed the variants which were already detected by VarScan (these are coloured red in the mutations list in Supplementary Table 2).

**Calculating the probability of parallel mutations in fusion genes:** The probability that the observed mutations affect fusion genes in all eight independently evolved lines can be conservatively calculated as follows. First, cheater morphotype evolved in all eight evolution lines independently with a number of mutations per strain of ~15 (conserved estimate based on Supplementary Table 2). Second, the probability of a mutation hitting a fusion gene will be 75/10000, given about ~75 known fusion genes and a total number of protein-coding genes in *N. crassa* to be around 10000. Putting these data together in a binomial test would yield a *P*-value of 3.991e-06: The R code line binom.test(8,8*15,75/10000). To correct this probability for the strains that did not evolve cheater morphotype the same test can be applied as follows. Binom.test(8,8*15,(10000-75)/10000) yielding a *P*-value < 2.2e-16. This negligible contribution would not affect the overall conclusion that the mutations hitting fusion genes eight times independently is not a chance effect.

**Competition assays:** To determine competitive success of fusion mutants against the wild type, we performed pairwise competition assays as follows. The strains were pre-grown in VMM slants for 6–7 days, and washed with sterile MQ-water to obtain spore suspensions. The spore concentrations were brought to 4∙10^7^ sp/ml and mixed at different ratios (from 10% to 90% of Δ*so* in the mixes with ~10% increments). We performed a separate experiment covering a range 5–40% of Δ*so* with ~5% increments to look at the competition dynamics with the Δ*so* frequency at around 25%. Both experiments included monoculture controls. 50 μl of the spore mixes were mat-inoculated in triplicates onto the surface of a slanted VMM medium, left horizontally to let the spore suspension to soak for 15 min, and then incubated upright for 4 days (or for 1–6 days to measure competitiveness in time, see section “No temporal benefit of the Δ*so*-cheater” in Supplementary Discussion). A small quantity of inoculum was spread onto 4–8 counting plates to estimate the realised initial frequency by phenotype counts, and correct for this value later when calculating competitiveness. After the competition, the spores were washed with 5 ml water (vortexed for ~20 seconds), diluted and inoculated onto counting plates. On counting plates, the fusion mutants could be distinguished by phenotype from the wild-type colonies (Supplementary Fig. 4), hence by counting each type the end frequency of the mutant could be determined. However, there is a possibility of errors due to stochastic effects on the phenotypes and a chance of chimera formations between the two genotypes masking the phenotype. To verify this phenotypic approach, we analysed the Δ*so*/wild type competitions (with 10% and 90% starting ratios of Δ*so*) using qPCR (Supplementary Fig. 1, see section “Frequency of Δ*so* cheater in heterokaryons”), which yielded conceptually similar results. The discrepancy at low starting frequency of Δ*so* may indicate underestimation of mutant counts due to its recessiveness in a heterokaryon state with the wild type.

For seven other fusion mutants (Δ*mak-1,* Δ*ham-3*, Δ*ham-4,* Δ*ham-5*, Δ*ham-6*, Δ*ham-7*, Δ*ham-8*), the competitions were done at a single starting frequency (~10%) and analysed by counting as with Δ*so*, since the mutants could be distinguished by phenotype on counting plates (Supplementary Fig. 4). The statistical significance of competitions was tested using one-sided *t*-test: Δ*so*/wt: *P*-value = 0.0142975; Δ*mak-1*/wt: *P*-value = 0.999931; Δ*ham-3*/wt: *P*-value = 0.994715; Δ*ham-4*/wt: *P*-value = 0.0323107; Δ*ham-5*/wt: *P*-value = 0.00537052; Δ*ham-6*/wt: *P*-value = 0.0145270; Δ*ham-7*/wt: *P*-value = 0.00190842; Δ*ham-8*/wt: *P*-value = 0.600992).

**Spore yield:** Spore yields were determined as in ^8^, except that the counting was done mostly with CASY TT Cell Counter (OMNI Life Science & Co KG, Germany). Occasionally though the haemocytometer was used. By counting random spore samples, we verified these two methods produce similar results.

**Heterokaryon formation:** To obtain heterokaryotic cultures, the *inl* and *pan* deficient strains were pre-grown separately in VMM (inositol and pantothenic acid supplemented) slants for 6–7 days, to generate sufficient number of spores. The spores were washed off with sterile MQ-water. The *inl* and *pan* deficient spores were mixed in 1:1 ratio and mat-inoculated on fresh VMM (no supplementation) slants to allow only heterokaryotic mycelium to propagate via markers complementation, as a result of the occasional fusion between the deficient strains. The emerged heterokaryotic colony allowed to sporulate (i.e. incubated for 4 days), spores were collected with sterile MQ-water and appropriate dilutions were plated on counting plates (no supplementation) to allow only heterokaryotic spores to germinate. The agar plug with a piece of a colony (~1 mm^3^) was used in subsequent experiments.

**Frequency of Δ*so* cheater in heterokaryons:** To determine Δ*so* cheater dynamics in heterokaryotic mycelium and derived spores, we isolated 10 independent heterokaryotic colonies (5 of wt^-inl^ + Δ*so*^-pan^ and 5 wt^-pan^ + Δ*so*^-inl^), which were randomly picked from non-supplemented counting plates (see section “Heterokaryon formation”), and let them grow in a ‘race tube’ (made of 50-ml disposable pipette, ~25 cm long) filled with ~1 cm wide VMM medium layer. After the mycelium has reached the end, the tubes were cut open, mycelium was collected from the start and the end of the tube and snap-frozen in liquid nitrogen and put to -80 °C for the subsequent DNA extraction (see section “DNA extraction”). Part of this mycelium was put into small glass tubes filled with non-slanted 0.5 ml of VMM. The reasoning behind using non-slanted small tubes was to restrict the mycelial outgrowth from the plug as much as possible. This way, the inoculated mycelium would mostly produce spores, limiting intra-mycelial allele dynamics upon mycelial outgrowth. After 7 days, the spores were collected with sterile MQ-water, centrifuged and stored at -80°C for the subsequent DNA extraction (see section “DNA extraction”). This experiment was performed both in VMM supplemented with inositol and pantothenic acid, and in non-supplemented VMM, to check how the heterokaryon enforcement would affect the cheater’s dynamics (see section “Heterokaryon enforcement when measuring Δ*so*-cheater frequency in mycelium and spores” in Supplementary Discussion).

qPCRs on gDNA were performed to determine the frequency of Δ*so* and wild-type nuclei within the heterokaryotic mycelium and derived spores. The principle was to differentially quantify the *hygB* gene (i.e. Δ*so* nuclei) and *so* gene (i.e. wild-type nuclei) in a gDNA sample derived from a heterokaryon between the Δ*so* cheater and the wild type. Each qPCR reaction (8 μl) was run in six technical replicates and per reaction contained: 4 μl 2x iQ SYBR Green SuperMix (Bio-Rad), 0.16 μl 10 uM of each primer, 3 μl of gDNA (18–990 pg), 0.68 μl sterile MQ-water. To amplify the *hygB* gene, we used hygB_2f/hygB_2r primer pair (5’->3’) CGGTTTCCACTATCGGCGAG and GGGCGTATATGCTCCGCATT. For the *so* gene, we used so_1f/so_1r primer pair (5’->3’) GTACCTCCTCTTCCAGCACC and CACCTTGGGGAACAGACCTT. The amplification program was set to 3 min 95°C, followed by 40 cycles of 30 sec at 95°C and 30 sec at 60°C. The reactions were run in a Bio-Rad CFX96 thermocycler. The qPCR results were analysed in Bio-Rad CFX Manager v.2.0 software. Baseline threshold line was set arbitrarily at the exponential phase of PCR (500 relative fluorescence units). Six technical replicates were averaged and Ct difference was calculated, based on which the frequency of each allele could be determined. Primer efficiencies and the overall method principle were verified in pilot qPCR runs on gDNA samples containing known ratios of each genotype.

For the statistics, we used linear regression and one-sided *t*-test to see whether the differences in Δ*so* dynamics between the end and the start of the mycelium and between the spores and mycelium are significant, with the following results:

##linear regression in NO SUPPL END/START MYCELIUM (F-statistic 1,8 = 0.5143, df=8, Multiple R-squared: 0.0604, *P*=0.4937);

##linear regression in YES SUPPL END/START MYCELIUM (F-statistic 1,8 = 5.309, df=8, Multiple R-squared: 0.3989, *P*=0.05014);

##one-sample one-tailed/one-sided (we are specifically interested if it is LESS than 1) *t*-test of NO SUPPL END/START MYCELIUM against 1 (mean = 0.8576329, t = -3.6702, df = 9, *P*-value = 0.002577**);

##one-sample one-tailed/one-sided (we are specifically interested if it is LESS than 1) *t*-test of YES SUPPL END/START MYCELIUM against 1 (mean = 0.6131703, t = -11.692, df = 9, *P*-value = 4.803e-07***);

##linear regression in NO SUPPL SPORES/MYCELIUM (F-statistic 1,18 = 52.97, df=18, Multiple R-squared: 0.7464, *P* =9.176e-07***);

##linear regression in YES SUPPL SPORES/MYCELIUM (F-statistic 1,18 = 15.71, df=18, Multiple R-squared: 0.466, *P* =0.0009106***).

**Competitions of Δ*so* against the vegetatively incompatible strains:** To see whether fusion is required for the cheating, the Δ*so* mutant (*het* Cde) was competed against vegetatively incompatible strains, including vegetatively compatible (*het* Cde) wild type strain as positive control. The vegetatively incompatible strains carried a different set of *het* genes (cDE, CdE, cde) which would block the successful fusion, as only the same suit of *het* alleles would allow this. The competitions were set up as explained before (see section “Competition assays”) at two starting ratios – 10% and 90%, in triplicates. After the competition, appropriate dilutions of the collected spores were plated counting plates for phenotyping, as well as frozen for the subsequent gDNA extraction qPCR analyses, as explained before (see section “Frequency of Δ*so* cheater in heterokaryons”). The results of the two methods (phenotyping and qPCR) were consistent (Supplementary Fig. 7); the results obtained with the qPCR are presented in the main text of this manuscript (Fig. 2b).

**Transfer experiments:** To track the dynamics of Δ*so* in the mixture with the wild type, the transfer experiments were performed as follows. The strains (*inl* or *pan* deficient) were pre-grown in VMM slants for 6–7 days, and the spores were collected with sterile MQ-water and brought to 4∙10^7^ sp/ml. The spores of Δ*so* were mixed with wild-type spores in 1:9 ratio and 50 μl of the mixes were mat-inoculated on *inl* and *pan* supplemented VMM slants, as before (see section “Competition assays”). After 4 days, the spores were washed with 5 ml MQ-water, the total spore yield was determined, 1% of the total spores (50 μl) were transferred to a fresh VMM (inositol and pantothenic acid supplemented) slant. At the same time, an appropriate dilution of those spores was inoculated on three types of counting plates (4 plates with inositol; 4 plates with pantothenic acid; 4 plates without supplementation) to determine the frequency of each genotype, including heterokaryotic spores. The process was repeated every 4 days for 4 transfers in triplicates. Δ*so*^-inl^/Δ*so*^-pan^ (1:1 starting ratio) transfers were done in parallel as a control to check for the degree of heterokaryon formation between Δ*so* and Δ*so*. To account for possible marker effects, the Δ*so* /wt transfer experiments were also done with reciprocal marker combinations.

**Statistical analyses and graphs:** Statistical analyses were performed in R 3.6.1^49^. The graphs were made using ggplot2 R package^50^ and aesthetics edited in Adobe Illustrator 2020.

**Data availability:** Raw sequencing data was deposited at NCBI BioProject number PRJNA605104. The web-link for reviewers: <https://dataview.ncbi.nlm.nih.gov/object/PRJNA605104?reviewer=s2h97d05ubpjqdemfgv2d4o20l>.

**Code availability:** Custom scripts for mutation analysis are available upon request.

2. **Supplementary Discussion**

**No temporal benefit of the Δ*so*-cheater:** An alternative hypothesis that could explain a competitive benefit of Δ*so* is that Δ*so*-rich mycelia sporulate earlier than wild type. The rationale for this idea is that sporulation is induced when local resources are exhausted, which happens earlier in a more fragmented fungal colony. We tested and rejected this hypothesis (Supplementary Fig. 2). First, Δ*so* was not overrepresented during the earlier stages of sporulation relative to its frequency later on. Second, we did not find a relative benefit of Δ*so* nuclei in competition with an incompatible wild-type strain (Fig. 2b, Supplementary Fig. 7).

**Frequency dependence of the benefit of the Δ*so*-cheater:** Our finding that the Δ*so* cheater has a benefit in the heterokaryon, explains its initial selection, since a nucleus with a *de novo* fusion mutation will find itself in a heterokaryon. At this initial low frequency, mutant nuclei will have an increased likelihood to end in the spores of this heterokaryon. With increasing frequency, increasing numbers of spores will be homokaryotic for the fusion deficiency. Upon germination, a fraction of these will be fused with wild-type or heterokaryotic mycelia, thus forming new heterokaryons, and the rest will develop into homokaryotic mycelia that have low reproductive success. Figure 1a shows that Δ*so* has a benefit below ~30% when starting from homokaryotic Δ*so* and wild-type spores, and Figure 3b that it has a benefit up to a frequency of ~60% when starting from a heterokaryon. The difference implies that in addition to frequency dependence within the heterokaryon, there is also frequency dependence of the chance to fuse (Fig. 4b, c). This can be understood by considering the probability that a fusion mutant fuses with at least one wild-type individual. At low frequencies, the Δ*so* mutant is mostly surrounded by wild type, maximizing its chance one of these fuses with it. With increasing frequencies of the Δ*so* the proportion of wild-type contacts decreases thus reducing the chance that at least one of those fuses with a focal Δ*so*. Moreover, wild-type colonies increasingly become fragmented due to Δ*so* patches, and more Δ*so*-rich heterokaryons increasingly segregate into homokaryotic sectors, reducing connectivity between supportive somatic and reproductive structures. This provides a relative benefit to wild type, restricts the opportunities of Δ*so* to end in spores upon fusion with a wild-type colony and explains further reduction in overall spore yield.

**Heterokaryon enforcement when measuring Δ*so*-cheater frequency in mycelium and spores:** Supplementing or not supplementing the media affects heterokaryon maintenance. Supplementing the medium would not enforce it, while not supplementing the media would enforce it, because of deficiencies complementation between genotypes. The reason behind including these treatments was to test if non-enforced conditions would disrupt the heterokaryon by potential outgrowth of either of the two genotypes. We did not see that, as we could still detect both genotypes present in a chimera. Yet, at non-enforced conditions, the drop in frequency of Δ*so* in the mycelium was stronger, than at enforced conditions (Supplementary Fig. 8a). However, we saw almost identical Δ*so*-cheater dynamics between the two treatments upon sporulation (Supplementary Fig. 8b). Since non-enforced conditions resemble the conditions of the evolution experiment, we present the data coming from the supplemented conditions in the main text (Fig. 3).

3. **Supplementary Figures**

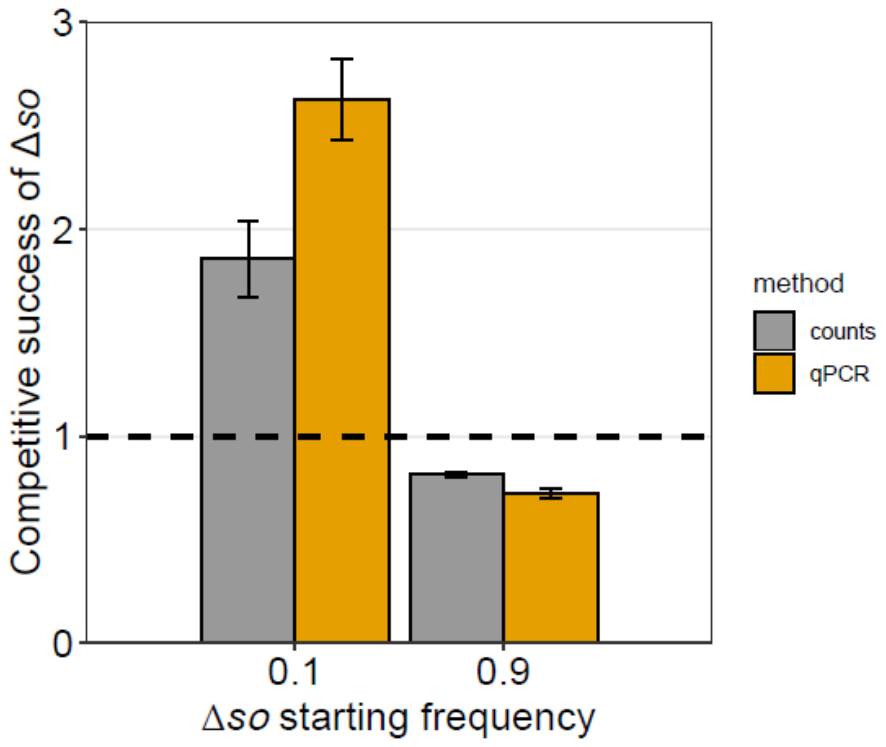

**Supplementary Figure 1.** **Competitive success of Δ*so* against the wild type at low (~10%) and high (~90%) frequency as measured by phenotype counts and by qPCR.** Error bars are 95% confidence intervals calculated from three biological replicates.

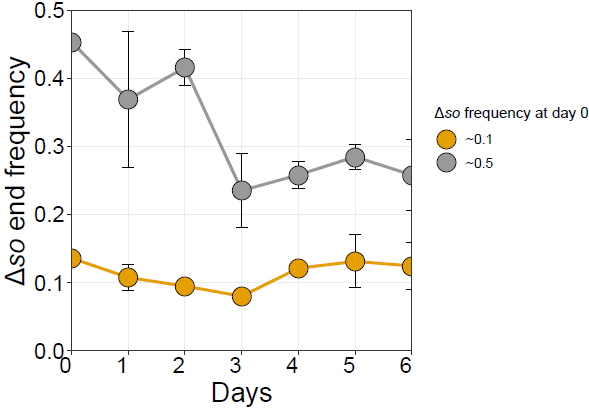

**Supplementary Figure 2.** **No temporal benefit of Δ*so* against wild type**. Error bars are 95% confidence intervals calculated from three biological replicates.

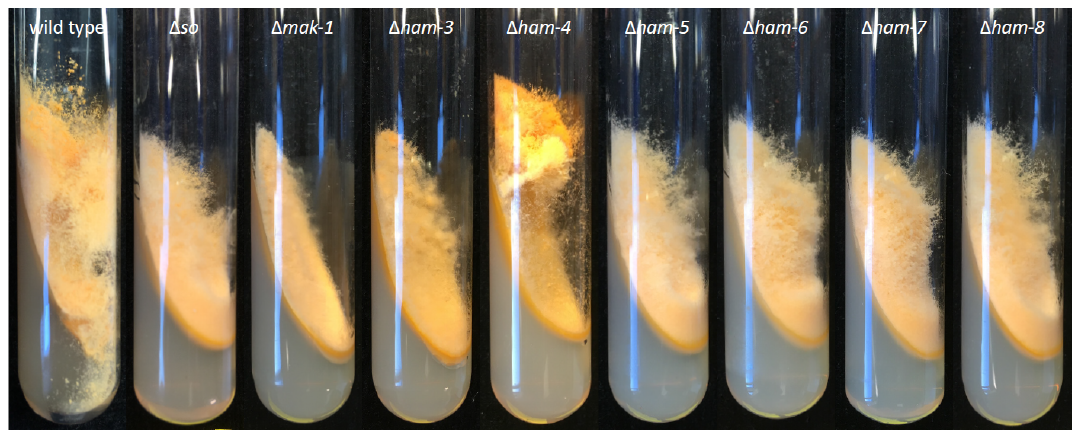

**Supplementary Figure 3.** **Phenotypes of fusion mutants (and the wild type, for contrast) used in the study.** 7-day old cultures on slanted VMM medium.

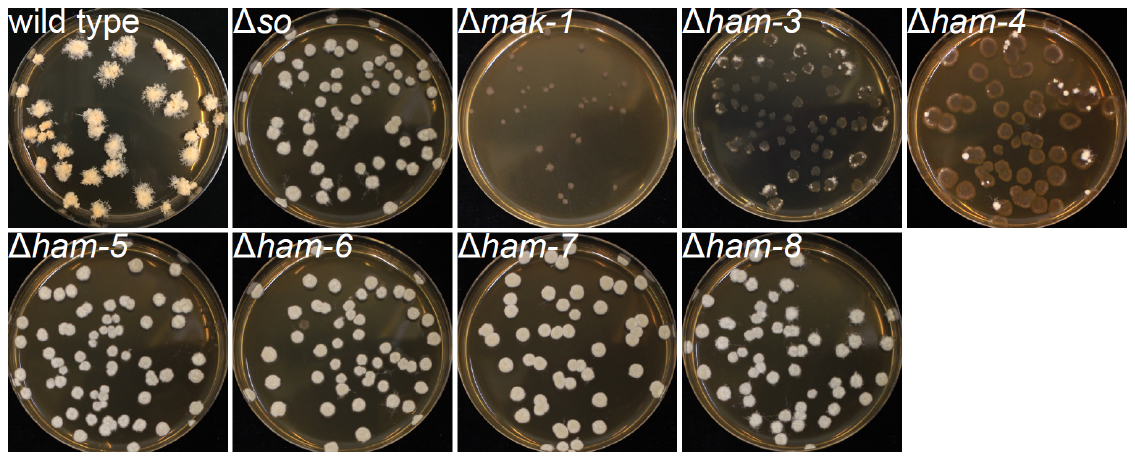

**Supplementary Figure 4.** **Phenotypes of fusion mutants (and the wild type, for contrast) used in the study.** 7-day old colonies on 9-cm Petri plates with sorbose-VMM (i.e. counting plates).

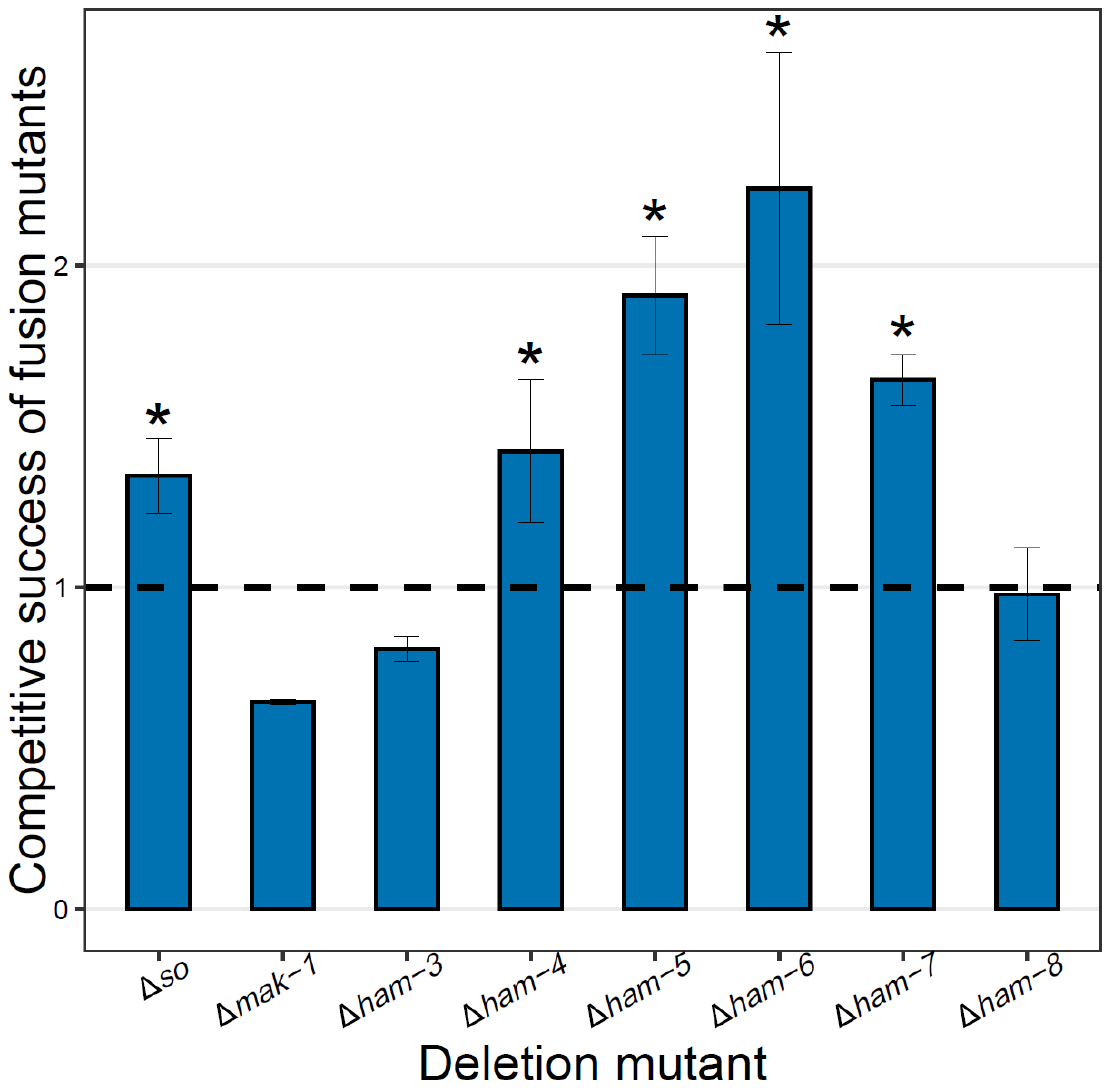

**Supplementary Figure 5.** **Competitive success of fusion mutants at low starting frequency (~10%) against the wild type.** Error bars are 95% confidence intervals calculated from three biological replicates. Asterisks indicate statistically significant higher competitiveness than wild type. For statistics, see section “Competition assays”.

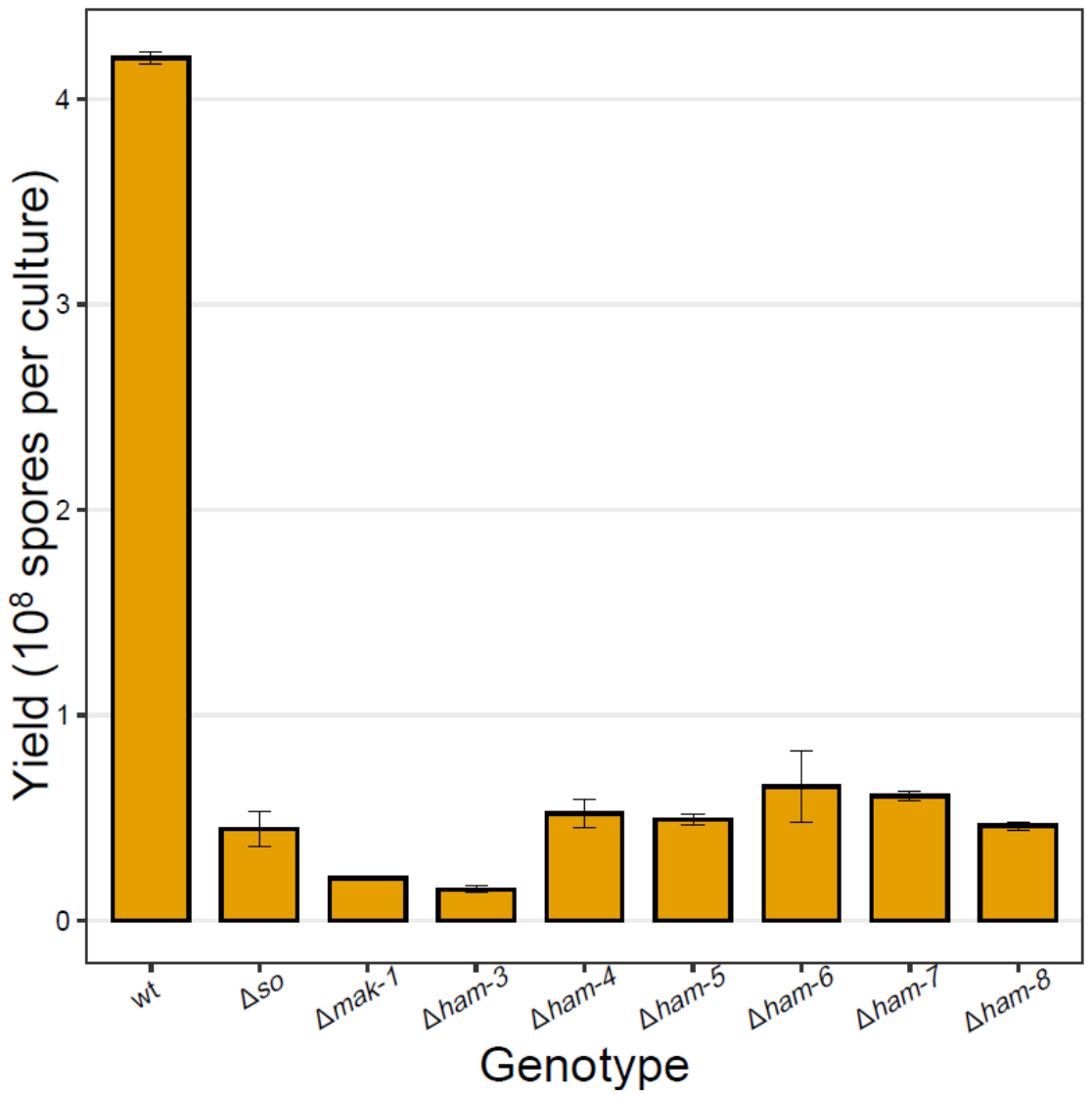

**Supplementary Figure 6.** **Spore yield of fusion mutants in 4-d-old VMM slants (along with the wild type control for contrast).** Error bars are 95% confidence intervals calculated from three biological replicates.

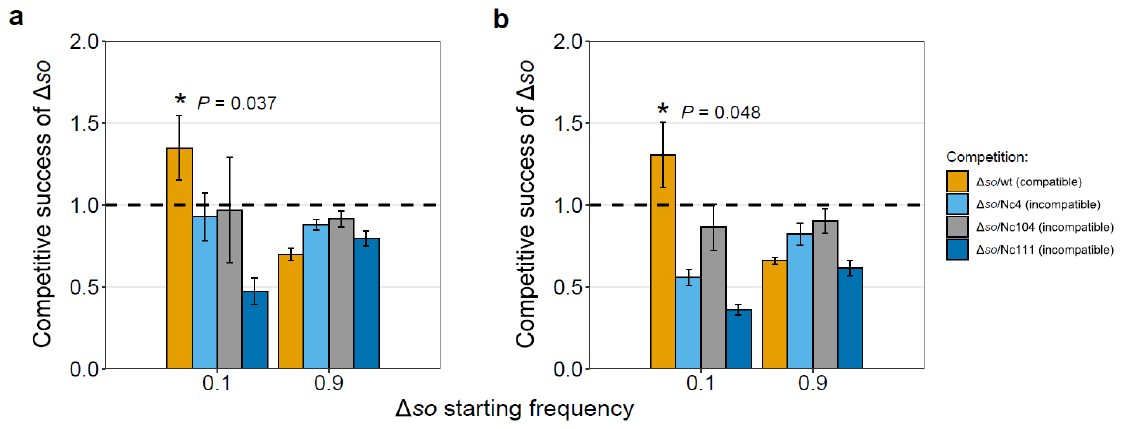

**Supplementary Figure 7.** **Competitive success of Δ*so* against vegetatively compatible (wt) and incompatible (Nc4/Nc104/Nc111) wild types at low (~10%) and high (~90%) frequencies.** **a,** As measured by phenotype counts (one-sided *t-*test, t-value = 3.469689, df = 2, *P* = 0.0369835). **b,** As measured by qPCR (one-sided *t-*test, t-value = 2.986799, df = 2, *P* = 0.0480968). Error bars are 95% confidence intervals calculated from three biological replicates.

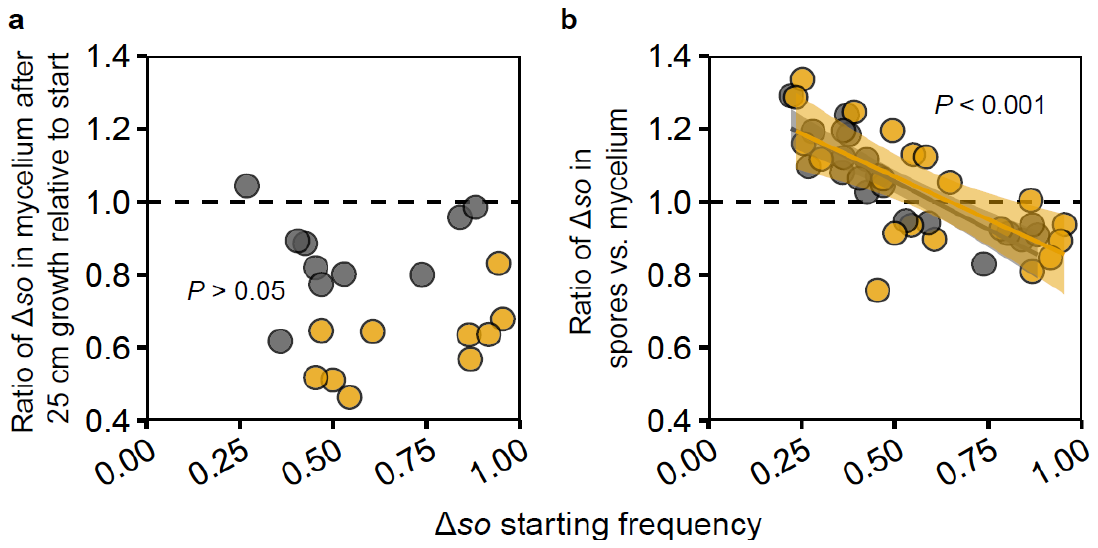

**Supplementary Figure 8.** **How Δ*so* realizes a competitive benefit in chimeras.** **a,** During mycelial growth, the frequency of Δ*so* decreased (one-sided *t-*test, *P* < 0.01), irrespectively of the starting frequency (linear regression, *P* > 0.05). A condition not enforcing the heterokaryon formation (orange dots) showed a stronger reduction of Δ*so* during somatic growth relative to strains that enforce heterokaryon formation (grey dots). **b,** In contrast, during spore formation Δ*so* nuclei have a benefit over wild-type nuclei as long as starting frequencies remain below ~60%, irrespective of whether heterokaryon formation is enforced (grey dots) or not (orange dots). Shaded areas around the trendlines are 95% confidence areas around the fitted lines. For details on statistics, see section “Frequency of Δ*so* cheater in heterokaryons”.

4. **Supplementary Tables**

| **Supplementary Table 1. Raw reads and mapping statistics of the sequenced *Neurospora crassa* morphotypes.** | | | | | | | |
| --- | --- | --- | --- | --- | --- | --- | --- |
| **Evolution line** | **Morphotype code** | **Raw reads (paired-end)** | **Trimmed reads (paired-end, length>=70,q>=30)** | **% trimmed** | **Mapped to reference genome (mapping q>=20, samtools flagstat: N = mapped - supplementary - secondary)** | **% mapped** | **Average coverage (only covered bases)** |
| ancestor to lines 9-16 | Nc152 | 16261518 | 12316616 | 24.26 | 12148931 | 98.64 | 41.6636 |
| ancestor to lines 1-8 | Nc159 | 16254600 | 12380134 | 23.84 | 12276868 | 99.17 | 41.3514 |
| 1 | 1t1 | 9685772 | 7413860 | 23.46 | 7356736 | 99.23 | 24.9313 |
|  | 1t2 | 8114430 | 6986582 | 13.90 | 6791561 | 97.21 | 23.8172 |
| 2 | 2t1 | 9345284 | 7616010 | 18.50 | 7547200 | 99.10 | 25.6872 |
| 3 | 3t1 | 8134722 | 7148050 | 12.13 | 7059575 | 98.76 | 24.6833 |
|  | 3t2 | 9729360 | 7819680 | 19.63 | 7742111 | 99.01 | 26.2718 |
| 4 | 4t1 | 8075234 | 7223682 | 10.55 | 7075664 | 97.95 | 25.0979 |
|  | 4t2 | 9706892 | 7407036 | 23.69 | 7359402 | 99.36 | 25.1834 |
|  | 4t3 | 8126006 | 7401912 | 8.91 | 7254109 | 98.00 | 25.7924 |
| 5 | 5t1 | 8131214 | 7324702 | 9.92 | 7227729 | 98.68 | 26.0144 |
|  | 5t2 | 8132348 | 7256980 | 10.76 | 7169275 | 98.79 | 26.4282 |
| 6 | 6t1 | 9717026 | 7457396 | 23.25 | 7404300 | 99.29 | 25.5318 |
|  | 6t2 | 8143338 | 7308602 | 10.25 | 7215427 | 98.73 | 25.7273 |
| 7 | 7t1 | 8142850 | 7318600 | 10.12 | 7216574 | 98.61 | 25.6734 |
|  | 7t2 | 8142338 | 7207082 | 11.49 | 7117221 | 98.75 | 26.2664 |
| 8 | 8t1 | 8134808 | 7350744 | 9.64 | 7257171 | 98.73 | 25.8312 |
|  | 8t2 | 8083878 | 7164486 | 11.37 | 7008072 | 97.82 | 24.7822 |
| 9 | 9t1 | 9762050 | 7679238 | 21.34 | 7614671 | 99.16 | 25.7705 |
|  | 9t2 | 9762066 | 7666062 | 21.47 | 7601877 | 99.16 | 25.789 |
| 10 | 10t1 | 9745086 | 7723500 | 20.74 | 7648961 | 99.03 | 25.9672 |
|  | 10t2 | 8138704 | 7358190 | 9.59 | 7264022 | 98.72 | 26.055 |
| 11 | 11t1 | 9604228 | 7439098 | 22.54 | 7380601 | 99.21 | 25.6529 |
|  | 11t2 | 9748028 | 7820730 | 19.77 | 7746580 | 99.05 | 26.347 |
|  | 11t3 | 9443604 | 7553618 | 20.01 | 7483980 | 99.08 | 25.3585 |
| 12 | 12t1 | 9769916 | 7449592 | 23.75 | 7395346 | 99.27 | 25.169 |
|  | 12t2 | 9237560 | 7027008 | 23.93 | 6973514 | 99.24 | 23.8072 |
| 13 | 13t1 | 9588674 | 7455312 | 22.25 | 7385870 | 99.07 | 24.9664 |
|  | 13t2 | 8136636 | 7340408 | 9.79 | 7254748 | 98.83 | 26.2485 |
|  | 13t3 | 9572632 | 7541784 | 21.22 | 7466937 | 99.01 | 25.1583 |
|  | 13t4 | 9741268 | 7545120 | 22.54 | 7471190 | 99.02 | 25.0438 |
| 14 | 14t1 | 9473952 | 7409758 | 21.79 | 7337594 | 99.03 | 24.7141 |
|  | 14t2 | 8831642 | 6933110 | 21.50 | 6863054 | 98.99 | 23.2501 |
| 15 | 15t1 | 9757916 | 7599888 | 22.12 | 7532955 | 99.12 | 25.5648 |
|  | 15t2 | 9721700 | 7291480 | 25.00 | 7240678 | 99.30 | 24.82 |
| 16 | 16t1 | 9520354 | 7047796 | 25.97 | 6994709 | 99.25 | 23.6181 |
|  | 16t2 | 9358704 | 7244788 | 22.59 | 7170883 | 98.98 | 23.909 |
|  | 16t3 | 9217438 | 6975760 | 24.32 | 6901601 | 98.94 | 21.9068 |

**Supplementary Table 2. The detected mutations in the evolved *Neurospora crassa* morphotypes relative to their respective ancestors.** Please see separate Supplementary_Table_2.xlsx file

| **Supplementary Table 3. Strains of *Neurospora crassa* used in the study.** | | | | |
| --- | --- | --- | --- | --- |
| **Genotype** | **Gene(s)# affected** | **Obtained from FGSC (#), or from a cross in this study, or a reference** | **Internal code** | ***het*** |
| *mat* A |  | 2489 | 147 | Cde |
| *mat* a |  | 4200 | 148 | Cde |
| *inl*; *mat* A | NCU06348 (*inl*) | ^8^ | 152 | Cde |
| *pan-2*; *mat* A | NCU10048 (*pan-2*) | ^8^ | 156 | Cde |
| *al-2*; *pan-1*; *mat* A | NCU00585 (*al-2*); NCU08661? (*pan-1/pan-4*) | 1425 | 004 | cDE |
| *inl*; *mat* A | NCU06348 (*inl*) | 538 | 104 | CdE |
| *al-2*; *pan-1*; *mat* A | NCU00585 (*al-2*); NCU08661? (*pan-1/pan-4*) | 2662 | 111 | cde |
| Δ*fl*::*Hyg^r^*; *mat* A | NCU08726 (*fl*) | Δ*fl*::*Hyg^r^*; *mat* a x mat A | 192 | Cde |
| Δ*fl*::*Hyg^r^*; *mat* a | NCU08726 (*fl*) | 11044 | 166 | Cde |
| Δ*so*::*Hyg^r^*; *mat* A | NCU02794 (*so*) | 11293 | 159=167 | Cde |
| Δ*so*::*Hyg^r^*; *mat* a | NCU02794 (*so*) | Δ*so*::*Hyg^r^*; *mat* A x mat a | 193 | Cde |
| Δ*so*::*Hyg^r^*; *inl*; *mat* A | NCU02794 (*so*); NCU06348 (*inl*) | Δ*so*::*Hyg^r^*; *mat* a x *inl*; *mat* A | 198 | Cde |
| Δ*so*::*Hyg^r^*; *pan-2*; *mat* A | NCU02794 (*so*); NCU10048 (*pan-2*) | Δ*so*::*Hyg^r^*; *mat* a x *pan-2*; *mat* A | 235 | Cde |
| Δ*mak-1*::*Hyg^r^*; *mat* A | NCU09842 (*mak-1*) | 11320 | 169 | Cde |
| Δ*ham-3*::*Hyg^r^*; *mat* A | NCU08741 (*ham-3*) | 11299 | 168 | Cde |
| Δ*ham-4*::*Hyg^r^*; *mat* A | NCU00528 (*ham-4*) | 12081 | 179 | Cde |
| Δ*ham-5*::*Hyg^r^*; *mat* A | NCU01789 (*ham-5*) | 15046 | 184 | Cde |
| Δ*ham-6*::*Hyg^r^*; *mat* A | NCU02767 (*ham-6*) | 16993 | 185 | Cde |
| Δ*ham-7*::*Hyg^r^*; *mat* A | NCU00881 (*ham-7*) | 13776 | 183 | Cde |
| Δ*ham-8*::*Hyg^r^*; *mat* A | NCU02811 (*ham-8*) | Δ*ham-8*::*Hyg^r^*; *mat* a x mat A | 224 | Cde |
| Δ*ham-8*::*Hyg^r^*; *mat* a | NCU02811 (*ham-8*) | 17225 | 186 | Cde |

**Supplementary References** **(continued from the main text)**

36.	Colot, H. V. *et al.* A high-throughput gene knockout procedure for Neurospora reveals functions for multiple transcription factors. *Proc. Natl. Acad. Sci. U. S. A.* **103**, 10352–10357 (2006).

37.	McCluskey, K., Wiest, A. & Plamann, M. The fungal genetics stock center: a repository for 50 years of fungal genetics research. *J. Biosci.* **35**, 119–126 (2010).

38.	Davis, R. H. & de Serres, F. J. Metabolism of Amino Acids and Amines Part A. *Methods Enzymol.* **17**, 79–143 (1970).

39.	Lichius, A. & Lord, K. M. Chemoattractive mechanisms in filamentous fungi. *Open Mycol. J.* **8**, 28–57 (2014).

40.	Rogers, S. O. & Bendich, A. J. Extraction of DNA from plant tissues. in *Plant Molecular Biology Manual* 1–10 (Kluwer Academic Publishers, Dordrecht, 1988).

41.	Kofler, R. *et al.* Popoolation: a toolbox for population genetic analysis of next generation sequencing data from pooled individuals. *PLoS One* **6**, e15925 (2011).

42.	Li, H. Aligning sequence reads, clone sequences and assembly contigs with BWA-MEM. (2013).

43.	Li, H. *et al.* The sequence alignment/map format and SAMtools. *Bioinformatics* **25**, 2078–2079 (2009).

44.	Li, H. A statistical framework for SNP calling, mutation discovery, association mapping and population genetical parameter estimation from sequencing data. *Bioinformatics* **27**, 2987–2993 (2011).

45.	Koboldt, D. C. *et al.* VarScan 2: somatic mutation and copy number alteration discovery in cancer by exome sequencing. *Genome Res.* **22**, 568–576 (2012).

46.	Knaus, B. J. & Grünwald, N. J. vcfr: a package to manipulate and visualize variant call format data in R. *Mol. Ecol. Resour.* **17**, 44–53 (2017).

47.	Robinson, J. T. *et al.* Integractive genomics viewer. *Nat. Biotechnol.* **29**, 24–26 (2011).

48.	Thorvaldsdóttir, H., Robinson, J. T. & Mesirov, J. P. Integrative Genomics Viewer (IGV): high-performance genomics data visualization and exploration. *Brief. Bioinform.* **14**, 178–192 (2013).

49.	R Core Team. R: A language and environment for statistical computing. R Foundation for Statistical Computing, Vienna, Austria. (2019).

50.	Wickham, H. *ggplot2: elegant graphics for data analysis*. (Springer, 2016).
