## Supplementary Table 2 for "Somatic deficiency causes reproductive parasitism in a fungus"

**Supplementary Table 2. The detected mutations in the evolved *Neurospora crassa* morphotypes relative to their respective ancestors.**

| Ancestor | Evolved morphotype | Scaffold | Position (reference) | Ancestor | Alternative | Frequency of alternative allele in ancestor | Frequency of alternative allele in evolved | Total coverage in ancestor | Total coverage in evolved | Coverage of alternative allele in ancestor | Coverage of alternative allele in evolved | Mutation type | Effect/Location | Codon change | Amino acid change | Gene name | Gene code (NCU#) | Gene orientation | Comment |
|---|---|---|---|---|---|---|---|---|---|---|---|---|---|---|---|---|---|---|---|
| Nc152 | 10t1 | Supercontig_12.1 | 2220105 | C | G | 0 | 1 | 46 | 43 | 0 | 43 | SNP | UTR | | | | NCU16352 | < | hypothetical protein |
| Nc152 | 10t1 | Supercontig_12.1 | 9214368 | A | ATGT | 0.020833333 | 0.875 | 48 | 24 | 1 | 21 | INS | inframe | | | | NCU02782 | < | hypothetical protein |
| Nc152 | 10t1 | Supercontig_12.2 | 224553 | A | G | 0 | 1 | 25 | 12 | 0 | 12 | SNP | intergenic | | | | | | |
| Nc152 | 10t1 | Supercontig_12.2 | 963760 | A | AC | 0 | 0.826086957 | 36 | 23 | 0 | 19 | INS | intergenic | | | | | | |
| Nc152 | 10t1 | Supercontig_12.2 | 3201402 | C | T | 0 | 1 | 53 | 14 | 0 | 14 | SNP | missense | GCC->GTC | A603V | cdp-3 | NCU01522 | > | chromodomain protein-3 |
| Nc152 | 10t1 | Supercontig_12.3 | 37945 | A | G | 0 | 1 | 38 | 34 | 0 | 34 | SNP | missense | GAC->GGC | D23G | mus-41 | NCU09516 | > | DNA repair protein rad-5 |
| Nc152 | 10t1 | Supercontig_12.5 | 431012 | T | A | 0 | 1 | 43 | 23 | 0 | 23 | SNP | intergenic | | | | | | |
| Nc152 | 10t1 | Supercontig_12.5 | 1832138 | G | A | 0 | 1 | 31 | 18 | 0 | 18 | SNP | intergenic | | | | | | |
| Nc152 | 10t1 | Supercontig_12.5 | 4540860 | G | T | 0 | 1 | 47 | 25 | 0 | 25 | SNP | nonsense | GAG->TAG | E333* | pod-2 | NCU09544 | > | polarity defective-2 (508aa) |
| Nc152 | 10t1 | Supercontig_12.6 | 1478586 | C | CAG | 0.140350877 | 0.96 | 57 | 25 | 8 | 24 | INS | UTR | | | | NCU05569 | > | hypothetical protein |
| Nc152 | 10t1 | Supercontig_12.6 | 3552665 | T | C | 0 | 1 | 43 | 33 | 0 | 33 | SNP | UTR | | | nca-3 | NCU05154 | > | calcium P-type ATPase |
| Nc152 | 10t2 | Supercontig_12.1 | 6222574 | G | C | 0 | 0.933333333 | 27 | 15 | 0 | 14 | SNP | intergenic | | | | | | |
| Nc152 | 10t2 | Supercontig_12.1 | 9113439 | G | T | 0 | 1 | 44 | 22 | 0 | 22 | SNP | splice site | | | so | NCU02794 | > | Fso1 |
| Nc152 | 10t2 | Supercontig_12.3 | 513139 | T | C | 0 | 1 | 33 | 21 | 0 | 21 | SNP | intergenic | | | | | | |
| Nc152 | 10t2 | Supercontig_12.5 | 5240447 | T | C | 0 | 1 | 25 | 32 | 0 | 32 | SNP | missense | ACC->GCC | T12A | ad-7 | NCU04216 | < | amidophosphoribosyltransferase |
| Nc152 | 10t2 | Supercontig_12.7 | 1763876 | G | T | 0 | 1 | 46 | 27 | 0 | 27 | SNP | missense | GAG->GAT | E296D | gtp-26 | NCU06111 | > | guanine triphosphate binding-26 |
| Nc152 | 11t2 | Supercontig_12.1 | 9028197 | C | T | 0 | 1 | 51 | 31 | 0 | 31 | SNP | missense | ACC->ATC | T696I | os-1 | NCU02815 | > | histidine kinase |
| Nc152 | 11t2 | Supercontig_12.2 | 2434392 | T | C | 0 | 1 | 78 | 27 | 0 | 27 | SNP | intronic | | | ham-5 | NCU01789 | > | Hyphal anastamosis-5 protein |
| Nc152 | 11t2 | Supercontig_12.3 | 1426615 | A | G | 0 | 1 | 58 | 20 | 0 | 20 | SNP | missense | ACC->GCC | T260A | | NCU06443 | > | hypothetical protein |
| Nc152 | 11t2 | Supercontig_12.5 | 130571 | A | G | 0 | 1 | 37 | 19 | 0 | 19 | SNP | intergenic | | | | | | |
| Nc152 | 11t2 | Supercontig_12.6 | 3671737 | A | G | 0 | 1 | 51 | 26 | 0 | 26 | SNP | intergenic | | | | | | |
| Nc152 | 11t3 | Supercontig_12.1 | 8947584 | C | T | 0 | 1 | 46 | 29 | 0 | 29 | SNP | intronic | | | cpn-4 | NCU02839 | > | T-complex protein 1 |
| Nc152 | 11t3 | Supercontig_12.4 | 539373 | G | A | 0 | 1 | 61 | 23 | 0 | 23 | SNP | missense | CCC->TCC | P21S | ada-6 | NCU04866 | < | All development-altered regulator ADA-6 |
| Nc152 | 11t3 | Supercontig_12.4 | 1615790 | G | T | 0 | 1 | 49 | 20 | 0 | 20 | SNP | UTR | | | alg-3/NA | NCU06552/NCU06 | >/< | dolichyl-P-Man:Man(5)GlcNAc(2)-PP-dolichyl mannosyltransferase/hypothetical protein |
| Nc152 | 11t3 | Supercontig_12.4 | 2417948 | C | CT | 0 | 0.869565217 | 47 | 23 | 0 | 20 | INS | intergenic | | | | | | |
| Nc152 | 11t3 | Supercontig_12.5 | 5847910 | T | C | 0 | 1 | 32 | 15 | 0 | 15 | SNP | UTR | | | crp-61 | NCU08963 | < | 60S ribosomal protein L30 |
| Nc152 | 11t3 | Supercontig_12.6 | 1478586 | C | CAG | 0.140350877 | 0.962962963 | 57 | 27 | 8 | 26 | INS | UTR | | | | NCU05569 | > | hypothetical protein |
| Nc152 | 11t3 | Supercontig_12.7 | 1768228 | G | T | 0 | 1 | 15 | 13 | 0 | 13 | SNP | intergenic | | | | | | |
| Nc152 | 11t3 | Supercontig_12.7 | 3353685 | C | A | 0 | 1 | 42 | 35 | 0 | 35 | SNP | synonymous | ACC->ACA | T1117T | stk-14 | NCU02202 | > | serine/threonine kinase IREI |
| Nc152 | 12t1 | Supercontig_12.1 | 1276672 | G | A | 0 | 0.892857143 | 58 | 28 | 0 | 25 | SNP | missense | AGC->AAC | S1710N | rpo-9 | NCU02103 | > | DNA-directed RNA polymerase II largest subunit |
| Nc152 | 12t1 | Supercontig_12.3 | 3976680 | A | C | 0 | 0.885714286 | 57 | 35 | 0 | 31 | SNP | intergenic | | | | | | |
| Nc152 | 12t1 | Supercontig_10.21 | 42922 | AG | A | 0 | 0.93554007 | 249 | 574 | 0 | 537 | DEL | UTR | | | | NCU16017 | > | hypothetical protein |
| Nc152 | 12t2 | Supercontig_12.1 | 4560125 | C | T | 0 | 1 | 39 | 19 | 0 | 19 | SNP | intergenic | | | | | | |
| Nc152 | 12t2 | Supercontig_12.1 | 9111005 | C | T | 0 | 1 | 50 | 26 | 0 | 26 | SNP | nonsense | CAA->TAA | Q322* | so | NCU02794 | > | Fso1 |
| Nc152 | 12t2 | Supercontig_12.3 | 4728872 | G | A | 0 | 1 | 78 | 26 | 0 | 26 | SNP | intergenic | | | | | | |
| Nc152 | 12t2 | Supercontig_12.5 | 2979088 | G | A | 0 | 1 | 34 | 25 | 0 | 25 | SNP | UTR | | | | NCU01127 | > | DUF636 domain-containing protein |
| Nc152 | 12t2 | Supercontig_12.5 | 4852924 | A | ATC | 0 | 0.931034483 | 48 | 29 | 0 | 27 | INS | frameshift | | | | NCU04105 | > | hypothetical protein |
| Nc152 | 12t2 | Supercontig_12.6 | 4029590 | G | T | 0 | 0.9 | 24 | 30 | 0 | 27 | SNP | intergenic | | | | | | |
| Nc152 | 12t2 | Supercontig_12.7 | 3539323 | G | A | 0 | 1 | 65 | 23 | 0 | 23 | SNP | UTR | | | | NCU08148 | < | H+/nucleoside cotransporter |
| Nc152 | 12t2 | Supercontig_12.7 | 3605645 | C | A | 0 | 1 | 56 | 22 | 0 | 22 | SNP | intergenic | | | | | | |
| Nc152 | 13t1 | Supercontig_12.3 | 3519065 | T | G | 0 | 1 | 12 | 16 | 0 | 16 | SNP | intergenic | | | | | | |
| Nc152 | 13t1 | Supercontig_12.7 | 1859328 | G | A | 0 | 0.972222222 | 32 | 36 | 0 | 35 | SNP | intergenic | | | | | | |
| Nc152 | 13t2 | Supercontig_12.1 | 3203007 | C | A | 0 | 1 | 47 | 28 | 0 | 28 | SNP | missense | CTC->ATC | L231I | | NCU02686 | > | hypothetical protein |
| Nc152 | 13t2 | Supercontig_12.1 | 4478322 | C | T | 0 | 1 | 62 | 36 | 0 | 36 | SNP | missense | GCC->ACC | A442T | | NCU03080 | < | hypothetical protein |
| Nc152 | 13t2 | Supercontig_12.2 | 3977901 | C | A | 0 | 1 | 34 | 24 | 0 | 24 | SNP | intergenic | | | | | | |
| Nc152 | 13t2 | Supercontig_12.3 | 229521 | G | C | 0 | 1 | 39 | 28 | 0 | 28 | SNP | intergenic | | | | | | |
| Nc152 | 13t2 | Supercontig_12.4 | 2126056 | CG | C | 0 | 0.857142857 | 51 | 28 | 0 | 24 | DEL | frameshift | | | | NCU07710 | < | vacuolar membrane-associated protein iml-1 |
| Nc152 | 13t2 | Supercontig_12.4 | 4347843 | C | T | 0 | 1 | 43 | 17 | 0 | 17 | SNP | UTR | | | ens-3 | NCU07330 | > | endosome-3 |
| Nc152 | 13t2 | Supercontig_12.5 | 3717245 | A | G | 0 | 1 | 53 | 26 | 0 | 26 | SNP | UTR | | | mind-2 | NCU01344 | < | hypothetical protein |
| Nc152 | 13t2 | Supercontig_12.5 | 4134120 | G | C | 0 | 1 | 68 | 21 | 0 | 21 | SNP | intergenic | | | | | | |
| Nc152 | 13t2 | Supercontig_12.6 | 276105 | C | A | 0 | 1 | 26 | 26 | 0 | 26 | SNP | missense | GAG->GAT | E821D | trm-6 | NCU07068 | < | K(+)/H(+) antiporter 1 |
| Nc152 | 13t2 | Supercontig_12.6 | 1196149 | C | A | 0 | 1 | 19 | 34 | 0 | 34 | SNP | intronic | | | cah-3 | NCU05653 | > | carbonic anhydrase-3 |
| Nc152 | 13t2 | Supercontig_12.6 | 1420246 | G | A | 0 | 1 | 49 | 27 | 0 | 27 | SNP | synonymous | GAC->GAT | D318D | | NCU05586 | < | malate/L-lactate dehydrogenase |
| Nc152 | 13t2 | Supercontig_12.6 | 1714718 | G | T | 0 | 1 | 30 | 29 | 0 | 29 | SNP | intergenic | | | | | | |
| Nc152 | 13t2 | Supercontig_12.6 | 3191989 | G | C | 0 | 1 | 70 | 28 | 0 | 28 | SNP | missense | CCA->GCA | P732A | chol-6 | NCU05985 | < | glycerol-3-phosphate O-acyltransferase |
| Nc152 | 13t2 | Supercontig_12.6 | 3537843 | G | T | 0 | 1 | 21 | 19 | 0 | 19 | SNP | intergenic | | | | | | |
| Nc152 | 13t2 | Supercontig_12.7 | 2769484 | T | TC | 0 | 0.967741935 | 50 | 31 | 0 | 30 | INS | frameshift | | | wc-1 | NCU02356 | > | white collar 1 protein |
| Nc152 | 13t3 | Supercontig_12.3 | 1579084 | C | G | 0 | 1 | 54 | 26 | 0 | 26 | SNP | synonymous | TCG->TCC | S388S | | NCU06484 | < | hypothetical protein |
| Nc152 | 13t3 | Supercontig_12.3 | 5000186 | G | A | 0 | 1 | 47 | 25 | 0 | 25 | SNP | UTR | | | | NCU08537 | < | hypothetical protein |
| Nc152 | 13t3 | Supercontig_12.4 | 765361 | T | A | 0 | 1 | 48 | 27 | 0 | 27 | SNP | intergenic | | | | | | |
| Nc152 | 13t3 | Supercontig_12.4 | 1040905 | C | T | 0 | 1 | 14 | 32 | 0 | 32 | SNP | intergenic | | | | | | |
| Nc152 | 13t3 | Supercontig_12.4 | 5294608 | C | T | 0 | 0.956521739 | 36 | 23 | 0 | 22 | SNP | splice site | | | | NCU09917 | < | hypothetical protein |
| Nc152 | 13t3 | Supercontig_12.5 | 1481241 | C | T | 0 | 1 | 59 | 24 | 0 | 24 | SNP | UTR | | | | NCU11109 | > | ORM1 |
| Nc152 | 13t3 | Supercontig_12.5 | 3818804 | G | T | 0 | 1 | 19 | 13 | 0 | 13 | SNP | missense | TTG->TTT | L639F | cdc16 | NCU01377 | > | anaphase-promoting complex subunit cut9 |
| Nc152 | 13t3 | Supercontig_12.5 | 4134120 | G | C | 0 | 1 | 68 | 21 | 0 | 21 | SNP | intergenic | | | | | | |
| Nc152 | 13t3 | Supercontig_12.6 | 276105 | C | A | 0 | 0.966666667 | 26 | 30 | 0 | 29 | SNP | missense | GAG->GAT | E821D | trm-6 | NCU07068 | < | K(+)/H(+) antiporter 1 |
| Nc152 | 13t3 | Supercontig_12.6 | 1196149 | C | A | 0 | 1 | 19 | 19 | 0 | 19 | SNP | intronic | | | cah-3 | NCU05653 | > | carbonic anhydrase-3 |
| Nc152 | 13t3 | Supercontig_12.6 | 1399474 | G | C | 0 | 1 | 62 | 32 | 0 | 32 | SNP | missense | TTC->TTG | F754L | abc-5 | NCU05591 | < | ABC transporter CDR4 |
| Nc152 | 13t3 | Supercontig_12.6 | 3628577 | G | T | 0 | 1 | 23 | 42 | 0 | 42 | SNP | missense | CGC->CTC | R2585L | gt4-1 | NCU05132 | > | hypothetical protein |
| Nc152 | 13t3 | Supercontig_12.6 | 3725398 | T | A | 0 | 1 | 56 | 22 | 0 | 22 | SNP | intergenic | | | | | | |
| Nc152 | 13t3 | Supercontig_12.7 | 489296 | T | A | 0 | 1 | 30 | 19 | 0 | 19 | SNP | intergenic | | | | | | |
| Nc152 | 13t3 | Supercontig_12.7 | 2569116 | A | T | 0 | 1 | 12 | 15 | 0 | 15 | SNP | UTR | | | gpl-3 | NCU02416 | > | glycerophospholipid-3 |
| Nc152 | 13t3 | Supercontig_12.7 | 2769484 | T | TC | 0 | 0.878787879 | 50 | 33 | 0 | 29 | INS | frameshift | | | wc-1 | NCU02356 | > | white collar 1 protein |
| Nc152 | 13t3 | Supercontig_12.7 | 3839516 | C | A | 0 | 1 | 34 | 19 | 0 | 19 | SNP | UTR | | | | NCU05791 | < | SOM1 protein |
| Nc152 | 13t4 | Supercontig_12.2 | 1646178 | G | T | 0 | 1 | 47 | 28 | 0 | 28 | SNP | synonymous | GGC->GGA | G359G | msp-29 | NCU05452 | < | splicing factor 3b |
| Nc152 | 13t4 | Supercontig_12.3 | 1579084 | C | G | 0 | 1 | 54 | 21 | 0 | 21 | SNP | synonymous | TCG->TCC | S388S | | NCU06484 | < | hypothetical protein |
| Nc152 | 13t4 | Supercontig_12.3 | 5000186 | G | A | 0 | 1 | 47 | 16 | 0 | 16 | SNP | UTR | | | | NCU08537 | < | hypothetical protein |
| Nc152 | 13t4 | Supercontig_12.4 | 765361 | T | A | 0 | 1 | 48 | 31 | 0 | 31 | SNP | intergenic | | | | | | |
| Nc152 | 13t4 | Supercontig_12.5 | 3818804 | G | T | 0 | 1 | 19 | 12 | 0 | 12 | SNP | missense | TTG->TTT | L639F | cdc16 | NCU01377 | > | anaphase-promoting complex subunit cut9 |
| Nc152 | 13t4 | Supercontig_12.5 | 4134120 | G | C | 0 | 1 | 68 | 33 | 0 | 33 | SNP | intergenic | | | | | | |
| Nc152 | 13t4 | Supercontig_12.6 | 276105 | C | A | 0 | 1 | 26 | 26 | 0 | 26 | SNP | missense | GAG->GAT | E821D | trm-6 | NCU07068 | < | K(+)/H(+) antiporter 1 |
| Nc152 | 13t4 | Supercontig_12.6 | 1196149 | C | A | 0 | 1 | 19 | 33 | 0 | 33 | SNP | intronic | | | cah-3 | NCU05653 | > | carbonic anhydrase-3 |
| Nc152 | 13t4 | Supercontig_12.6 | 1399474 | G | C | 0 | 1 | 62 | 38 | 0 | 38 | SNP | missense | TTC->TTG | F754L | abc-5 | NCU05591 | < | ABC transporter CDR4 |
| Nc152 | 13t4 | Supercontig_12.6 | 3725398 | T | A | 0 | 1 | 56 | 20 | 0 | 20 | SNP | intergenic | | | | | | |
| Nc152 | 13t4 | Supercontig_12.7 | 489296 | T | A | 0 | 1 | 30 | 24 | 0 | 24 | SNP | intergenic | | | | | | |
| Nc152 | 13t4 | Supercontig_12.7 | 2569116 | A | T | 0 | 1 | 12 | 16 | 0 | 16 | SNP | UTR | | | gpl-3 | NCU02416 | > | glycerophospholipid-3 |
| Nc152 | 13t4 | Supercontig_12.7 | 2769484 | T | TC | 0 | 0.857142857 | 50 | 21 | 0 | 18 | INS | frameshift | | | wc-1 | NCU02356 | > | white collar 1 protein |
| Nc152 | 14t2 | Supercontig_12.1 | 6774836 | G | A | 0 | 1 | 44 | 27 | 0 | 27 | SNP | intergenic | | | | | | |
| Nc152 | 14t2 | Supercontig_12.2 | 1386139 | G | A | 0 | 1 | 39 | 19 | 0 | 19 | SNP | missense | GCC->ACC | A50T | | NCU05345 | > | zearalenone lactonase |
| Nc152 | 14t2 | Supercontig_12.3 | 3994980 | C | T | 0 | 1 | 69 | 20 | 0 | 20 | SNP | missense | GGT->AGT | G393S | acon-2 | NCU00478 | < | aconidiate-2 |
| Nc152 | 14t2 | Supercontig_12.4 | 353440 | G | T | 0 | 1 | 29 | 31 | 0 | 31 | SNP | intergenic | | | | | | |
| Nc152 | 14t2 | Supercontig_12.6 | 1478586 | C | CAG | 0.140350877 | 1 | 57 | 18 | 8 | 18 | INS | UTR | | | | NCU05569 | > | hypothetical protein |
| Nc152 | 14t2 | Supercontig_12.6 | 3537827 | T | TG | 0 | 0.875 | 22 | 24 | 0 | 21 | INS | intergenic | | | | | | |
| Nc152 | 14t2 | Supercontig_12.7 | 4191540 | C | T | 0 | 1 | 40 | 25 | 0 | 25 | SNP | intergenic | | | | | | |
| Nc152 | 15t1 | Supercontig_12.1 | 1170817 | A | C | 0 | 1 | 28 | 27 | 0 | 27 | SNP | missense | GAT->GCT | D172A | | NCU02122 | > | HCO3 |
| Nc152 | 15t1 | Supercontig_12.3 | 1302431 | C | G | 0 | 1 | 37 | 21 | 0 | 21 | SNP | missense | AAG->AAC | K506N | rbg-51 | NCU06409 | < | ribosome biogenesis-51 |
| Nc152 | 15t1 | Supercontig_12.4 | 3940479 | A | G | 0 | 1 | 57 | 22 | 0 | 22 | SNP | synonymous | CAA->CAG | Q131Q | gid-1 | NCU08021 | > | GPI inositol-deacylase |
| Nc152 | 15t1 | Supercontig_12.4 | 5391915 | T | A | 0 | 1 | 39 | 15 | 0 | 15 | SNP | intergenic | | | | | | |
| Nc152 | 15t1 | Supercontig_12.5 | 2896946 | C | T | 0 | 1 | 39 | 29 | 0 | 29 | SNP | intergenic | | | | | | |
| Nc152 | 15t2 | Supercontig_12.1 | 2418541 | C | A | 0 | 1 | 55 | 19 | 0 | 19 | SNP | missense | GAC->GAA | D365E | gsl-20 | NCU02468 | > | longevity-assurance protein |
| Nc152 | 15t2 | Supercontig_12.1 | 7456068 | C | T | 0 | 1 | 45 | 37 | 0 | 37 | SNP | intergenic | | | | | | |
| Nc152 | 15t2 | Supercontig_12.1 | 8876590 | G | T | 0 | 1 | 31 | 19 | 0 | 19 | SNP | missense | GAC->GAA | D163E | | NCU16483 | < | hypothetical protein |
| Nc152 | 15t2 | Supercontig_12.3 | 3641295 | C | A | 0 | 1 | 54 | 24 | 0 | 24 | SNP | UTR | | | ahd-2 | NCU00378 | > | alhehyde dehydrogenase-2 |
| Nc152 | 15t2 | Supercontig_12.5 | 5566006 | G | A | 0 | 1 | 65 | 24 | 0 | 24 | SNP | missense | CCA->TCA | P159S | | NCU04305 | < | hypothetical protein |
| Nc152 | 16t1 | Supercontig_10.21 | 5833 | A | C | 0 | 1 | 130 | 22 | 0 | 22 | SNP | intergenic | | | | | | |
| Nc152 | 16t1 | Supercontig_10.21 | 42922 | A | AG | 0 | 0.947692308 | 249 | 325 | 0 | 308 | INS | UTR | | | | NCU16017 | > | hypothetical protein |
| Nc152 | 16t2 | Supercontig_12.1 | 5226269 | A | G | 0 | 1 | 48 | 28 | 0 | 28 | SNP | intergenic | | | | | | |
| Nc152 | 16t2 | Supercontig_12.1 | 7873983 | C | CA | 0 | 0.80952381 | 42 | 21 | 0 | 17 | INS | UTR | | | rho-3 | NCU00600 | < | Small Rho-type GTPase |
| Nc152 | 16t2 | Supercontig_12.1 | 9042601 | C | T | 0 | 1 | 41 | 33 | 0 | 33 | SNP | nonsense | TGG->TAG | W277* | ham-8 | NCU02811 | < | hyphal anastamosis-8 protein |
| Nc152 | 16t2 | Supercontig_12.3 | 4772629 | G | A | 0 | 1 | 47 | 27 | 0 | 27 | SNP | intergenic | | | | | | |
| Nc152 | 16t2 | Supercontig_12.4 | 3838407 | C | G | 0 | 1 | 50 | 27 | 0 | 27 | SNP | missense | AAC->AAG | N30K | | NCU07994 | > | hypothetical protein |
| Nc152 | 16t2 | Supercontig_10.21 | 5833 | A | C | 0 | 1 | 130 | 21 | 0 | 21 | SNP | intergenic | | | | | | |
| Nc152 | 16t2 | Supercontig_10.21 | 42922 | A | AG | 0 | 0.894348894 | 249 | 407 | 0 | 364 | INS | UTR | | | | NCU16017 | > | hypothetical protein |
| Nc152 | 16t3 | Supercontig_12.1 | 1235164 | A | G | 0 | 1 | 34 | 19 | 0 | 19 | SNP | missense | ACC->GCC | T343A | div-8 | NCU02114 | > | G1/S-specific cyclin Cln1 |
| Nc152 | 16t3 | Supercontig_12.1 | 4732250 | TACA | T | 0 | 0.875 | 67 | 24 | 0 | 21 | DEL | splice site | | | | NCU03151 | < | Peroxisomal membrane protein |
| Nc152 | 16t3 | Supercontig_12.1 | 8483817 | C | A | 0 | 1 | 67 | 17 | 0 | 17 | SNP | intergenic | | | | | | |
| Nc152 | 16t3 | Supercontig_12.5 | 4897027 | C | T | 0 | 1 | 65 | 21 | 0 | 21 | SNP | UTR | | | | NCU04114 | > | hypothetical protein |
| Nc152 | 16t3 | Supercontig_12.7 | 3546055 | C | A | 0 | 1 | 78 | 27 | 0 | 27 | SNP | intergenic | | | | | | |
| Nc152 | 16t3 | Supercontig_10.21 | 5833 | A | C | 0 | 1 | 130 | 16 | 0 | 16 | SNP | intergenic | | | | | | |
| Nc152 | 16t3 | Supercontig_10.21 | 42922 | A | AG | 0 | 0.953020134 | 249 | 298 | 0 | 284 | INS | UTR | | | | NCU16017 | > | hypothetical protein |
| Nc152 | 9t1 | Supercontig_12.5 | 2706204 | A | G | 0 | 0.888888889 | 66 | 27 | 0 | 24 | SNP | UTR | | | mrp-27 | NCU01023 | < | 50S ribosomal protein L17 |
| Nc152 | 9t2 | Supercontig_12.1 | 3336649 | T | TG | 0 | 0.848484848 | 55 | 33 | 0 | 28 | INS | frameshift | | | | NCU02725 | < | hypothetical protein |
| Nc152 | 9t2 | Supercontig_12.1 | 9112515 | T | C | 0 | 1 | 58 | 27 | 0 | 27 | SNP | missense | CTG->CCG | L825P | so | NCU02794 | > | Fso1 |
| Nc152 | 9t2 | Supercontig_12.1 | 9398463 | CT | C | 0 | 1 | 19 | 14 | 0 | 14 | DEL | intergenic | | | | | | |
| Nc152 | 9t2 | Supercontig_12.2 | 3772672 | CT | C | 0 | 0.931034483 | 41 | 29 | 0 | 27 | DEL | intergenic | | | | | | |
| Nc152 | 9t2 | Supercontig_12.3 | 4636729 | A | G | 0 | 1 | 63 | 26 | 0 | 26 | SNP | intergenic | | | | | | |
| Nc152 | 9t2 | Supercontig_12.4 | 4401253 | C | A | 0 | 1 | 21 | 21 | 0 | 21 | SNP | intergenic | | | | | | |
| Nc152 | 9t2 | Supercontig_12.7 | 1113217 | A | G | 0 | 1 | 28 | 25 | 0 | 25 | SNP | intergenic | | | | | | |
| Nc159 | 1t1 | Supercontig_12.1 | 8374395 | T | C | 0 | 1 | 25 | 14 | 0 | 14 | SNP | intergenic | | | | | | |
| Nc159 | 1t1 | Supercontig_12.3 | 3865796 | G | T | 0 | 1 | 46 | 28 | 0 | 28 | SNP | missense | TGC->TTC | C136F | gnb-1 | NCU00440 | > | guanine nucleotide-binding protein beta subunit |
| Nc159 | 1t1 | Supercontig_12.5 | 4718894 | G | A | 0 | 1 | 37 | 20 | 0 | 20 | SNP | missense | GGT->AGT | G15S | gtp-19 | NCU01444 | > | rheb small monomeric GTPase RhbA |
| Nc159 | 1t2 | Supercontig_12.1 | 3092350 | C | G | 0 | 1 | 46 | 20 | 0 | 20 | SNP | UTR | | | eth-1 | NCU02657 | < | S-adenosylmethionine synthetase |
| Nc159 | 1t2 | Supercontig_12.1 | 3843079 | G | C | 0 | 1 | 13 | 12 | 0 | 12 | SNP | intergenic | | | | | | |
| Nc159 | 1t2 | Supercontig_12.1 | 9027185 | A | G | 0 | 1 | 40 | 23 | 0 | 23 | SNP | missense | ATT->GTT | I398V | os-1 | NCU02815 | > | histidine kinase |
| Nc159 | 1t2 | Supercontig_12.2 | 218411 | G | T | 0 | 1 | 47 | 19 | 0 | 19 | SNP | intergenic | | | | | | |
| Nc159 | 1t2 | Supercontig_12.3 | 1138309 | G | A | 0 | 1 | 48 | 28 | 0 | 28 | SNP | intergenic | | | | | | |
| Nc159 | 1t2 | Supercontig_12.3 | 3865796 | G | T | 0 | 1 | 46 | 31 | 0 | 31 | SNP | missense | TGC->TTC | C136F | gnb-1 | NCU00440 | > | guanine nucleotide-binding protein beta subunit |
| Nc159 | 1t2 | Supercontig_12.4 | 4345737 | T | C | 0 | 1 | 32 | 16 | 0 | 16 | SNP | UTR | | | NA/rbg-5 | NCU07332/NCU07 | >/< | hypothetical protein/ribosome biogenesis-56 |
| Nc159 | 1t2 | Supercontig_12.4 | 5525042 | A | G | 0 | 1 | 44 | 34 | 0 | 34 | SNP | intergenic | | | | | | |
| Nc159 | 1t2 | Supercontig_12.6 | 3723570 | G | T | 0 | 1 | 48 | 23 | 0 | 23 | SNP | intergenic | | | | | | |
| Nc159 | 1t2 | Supercontig_12.7 | 887735 | G | T | 0 | 1 | 33 | 21 | 0 | 21 | SNP | intergenic | | | | | | |
| Nc159 | 1t2 | Supercontig_12.9 | 96475 | C | T | 0 | 1 | 16 | 12 | 0 | 12 | SNP | intergenic | | | | | | |
| Nc159 | 2t1 | Supercontig_12.1 | 9027560 | A | G | 0 | 1 | 33 | 28 | 0 | 28 | SNP | missense | ACC->GCC | T523A | os-1 | NCU02815 | > | histidine kinase |
| Nc159 | 2t1 | Supercontig_12.2 | 1237019 | C | A | 0 | 1 | 17 | 10 | 0 | 10 | SNP | intergenic | | | | | | |
| Nc159 | 2t1 | Supercontig_12.2 | 1237021 | A | T | 0 | 1 | 16 | 11 | 0 | 11 | SNP | intergenic | | | | | | |
| Nc159 | 2t1 | Supercontig_12.3 | 625988 | C | T | 0 | 1 | 49 | 41 | 0 | 41 | SNP | synonymous | ACG->ACA | T260T | csc-1 | NCU05720 | < | chitin synthase export chaperone |
| Nc159 | 2t1 | Supercontig_12.3 | 2843487 | C | T | 0 | 1 | 63 | 21 | 0 | 21 | SNP | intergenic | | | | | | |
| Nc159 | 2t1 | Supercontig_12.5 | 996862 | T | G | 0 | 1 | 10 | 11 | 0 | 11 | SNP | intergenic | | | | | | |
| Nc159 | 2t1 | Supercontig_12.5 | 2924016 | G | A | 0 | 1 | 29 | 23 | 0 | 23 | SNP | missense | GAG->AAG | E55K | cah-1 | NCU01103 | > | carbonic anhydrase-1 |
| Nc159 | 2t1 | Supercontig_12.7 | 578351 | C | A | 0 | 1 | 44 | 28 | 0 | 28 | SNP | UTR | | | | NCU06867 | > | hypothetical protein |
| Nc159 | 3t1 | Supercontig_12.1 | 3966681 | A | C | 0 | 1 | 16 | 16 | 0 | 16 | SNP | intergenic | | | | | | |
| Nc159 | 3t1 | Supercontig_12.4 | 2120814 | G | A | 0 | 1 | 42 | 26 | 0 | 26 | SNP | nonsense | CAA->TAA | Q1771* | | NCU07710 | < | vacuolar membrane-associated protein iml-1 |
| Nc159 | 3t1 | Supercontig_12.7 | 2569116 | A | T | 0 | 1 | 22 | 19 | 0 | 19 | SNP | UTR | | | gpl-3 | NCU02416 | > | glycerophospholipid-3 |
| Nc159 | 3t1 | Supercontig_12.7 | 4108316 | G | T | 0 | 1 | 44 | 28 | 0 | 28 | SNP | missense | CCG->CAG | P112Q | | NCU05866 | < | hypothetical protein |
| Nc159 | 3t2 | Supercontig_12.2 | 1187565 | A | G | 0 | 1 | 13 | 26 | 0 | 26 | SNP | intergenic | | | | | | |
| Nc159 | 3t2 | Supercontig_12.2 | 1624027 | A | G | 0 | 1 | 48 | 31 | 0 | 31 | SNP | missense | CTT->CCT | L132P | | NCU05435 | < | hypothetical protein |
| Nc159 | 3t2 | Supercontig_12.2 | 4435674 | A | G | 0 | 1 | 30 | 28 | 0 | 28 | SNP | missense | GAG->GGG | E252G | gh13-3 | NCU09486 | > | glucan 1,4-alpha-maltohexaosidase |
| Nc159 | 3t2 | Supercontig_12.3 | 3008526 | C | A | 0 | 1 | 47 | 26 | 0 | 26 | SNP | intergenic | | | | | | |
| Nc159 | 3t2 | Supercontig_12.4 | 3090177 | G | A | 0 | 1 | 27 | 16 | 0 | 16 | SNP | intergenic | | | | | | |
| Nc159 | 3t2 | Supercontig_12.5 | 4854188 | C | A | 0 | 1 | 42 | 31 | 0 | 31 | SNP | missense | GCA->GAA | A988E | | NCU04105 | > | hypothetical protein |
| Nc159 | 3t2 | Supercontig_12.5 | 6241311 | C | G | 0 | 1 | 44 | 22 | 0 | 22 | SNP | intergenic | | | | | | |
| Nc159 | 3t2 | Supercontig_12.6 | 543575 | C | G | 0 | 0.833333333 | 34 | 12 | 0 | 10 | SNP | UTR | | | | NCU04721 | < | neutral ceramidase |
| Nc159 | 3t2 | Supercontig_12.6 | 2902490 | T | A | 0 | 1 | 16 | 10 | 0 | 10 | SNP | intergenic | | | | | | |
| Nc159 | 3t2 | Supercontig_12.7 | 2452592 | T | G | 0 | 1 | 17 | 10 | 0 | 10 | SNP | intergenic | | | | | | |
| Nc159 | 3t2 | Supercontig_12.8 | 155873 | A | C | 0 | 0.909090909 | 13 | 11 | 0 | 10 | SNP | intergenic | | | | | | |
| Nc159 | 4t1 | Supercontig_12.1 | 2929547 | G | A | 0 | 1 | 51 | 28 | 0 | 28 | SNP | missense | GGT->AGT | G132S | | NCU02617 | > | kelch repeat-containing protein |
| Nc159 | 4t1 | Supercontig_12.1 | 3437626 | AAC | A | 0 | 0.947368421 | 47 | 19 | 0 | 18 | DEL | UTR | | | znf-47 | NCU02752 | < | zinc finger transcription factor-47 |
| Nc159 | 4t1 | Supercontig_12.4 | 2124773 | A | T | 0 | 1 | 59 | 27 | 0 | 27 | SNP | nonsense | TTG->TAG | L451* | | NCU07710 | < | vacuolar membrane-associated protein iml-1 |
| Nc159 | 4t1 | Supercontig_12.5 | 3033662 | A | G | 0 | 1 | 40 | 21 | 0 | 21 | SNP | synonymous | CAA->CAG | Q591Q | | NCU01150 | > | UBA domain-containing protein 3 |
| Nc159 | 4t1 | Supercontig_12.5 | 4217867 | G | A | 0 | 1 | 41 | 34 | 0 | 34 | SNP | UTR | | | | NCU06674 | > | hypothetical protein |
| Nc159 | 4t2 | Supercontig_12.1 | 5600396 | G | A | 0 | 1 | 51 | 23 | 0 | 23 | SNP | UTR | | | | NCU08369 | < | rhodopsin family protein |
| Nc159 | 4t2 | Supercontig_12.1 | 7945776 | C | A | 0 | 1 | 52 | 22 | 0 | 22 | SNP | missense | TTC->TTA | F970L | osb-6 | NCU00579 | > | oxysterol-binding protein-6 |
| Nc159 | 4t2 | Supercontig_12.1 | 9027259 | C | G | 0 | 1 | 42 | 19 | 0 | 19 | SNP | missense | ATC->ATG | I422M | os-1 | NCU02815 | > | histidine kinase |
| Nc159 | 4t2 | Supercontig_12.2 | 2425215 | T | C | 0.025641026 | 1 | 39 | 32 | 1 | 32 | SNP | intergenic | | | | | | |
| Nc159 | 4t2 | Supercontig_12.3 | 4925793 | C | T | 0 | 0.826086957 | 60 | 23 | 0 | 19 | SNP | intergenic | | | | | | |
| Nc159 | 4t2 | Supercontig_12.6 | 3796126 | G | T | 0 | 0.952380952 | 29 | 21 | 0 | 20 | SNP | UTR | | | | NCU05085 | < | hypothetical protein |
| Nc159 | 4t3 | Supercontig_12.1 | 3611217 | G | T | 0 | 1 | 11 | 15 | 0 | 15 | SNP | intergenic | | | | | | |
| Nc159 | 4t3 | Supercontig_12.1 | 3887678 | C | G | 0 | 1 | 16 | 11 | 0 | 11 | SNP | intergenic | | | | | | |
| Nc159 | 4t3 | Supercontig_12.1 | 6889145 | G | A | 0 | 1 | 38 | 31 | 0 | 31 | SNP | intergenic | | | | | | |
| Nc159 | 4t3 | Supercontig_12.2 | 2425215 | T | C | 0.025641026 | 1 | 39 | 23 | 1 | 23 | SNP | intergenic | | | | | | |
| Nc159 | 4t3 | Supercontig_12.2 | 3559126 | C | A | 0 | 1 | 61 | 28 | 0 | 28 | SNP | missense | GGT->TGT | G226C | bud6 | NCU08468 | < | actin-interacting protein |
| Nc159 | 4t3 | Supercontig_12.3 | 3994646 | A | G | 0 | 1 | 34 | 27 | 0 | 27 | SNP | missense | ATC->ACC | I504T | acon-2 | NCU00478 | < | aconidiate-2 |
| Nc159 | 4t3 | Supercontig_12.3 | 3994649 | A | T | 0 | 1 | 35 | 27 | 0 | 27 | SNP | missense | ATA->AAA | I503K | acon-2 | NCU00478 | < | aconidiate-2 |
| Nc159 | 4t3 | Supercontig_12.5 | 3361351 | C | A | 0 | 1 | 47 | 32 | 0 | 32 | SNP | missense | CTT->ATT | L304I | coh-3 | NCU01247 | > | hypothetical protein |
| Nc159 | 4t3 | Supercontig_12.5 | 5145592 | A | G | 0 | 1 | 47 | 34 | 0 | 34 | SNP | synonymous | ACT->ACC | T43T | cot-2 | NCU04189 | < | glucan synthesis regulatory protein |
| Nc159 | 4t3 | Supercontig_12.7 | 833582 | C | A | 0 | 1 | 43 | 24 | 0 | 24 | SNP | UTR | | | | NCU06937 | > | hypothetical protein |
| Nc159 | 4t3 | Supercontig_12.7 | 2930528 | G | C | 0 | 1 | 30 | 11 | 0 | 11 | SNP | intergenic | | | | | | |
| Nc159 | 4t3 | Supercontig_12.7 | 3232948 | G | C | 0 | 0.978723404 | 50 | 47 | 0 | 46 | SNP | intronic | | | nup-10 | NCU02236 | > | hypothetical protein |
| Nc159 | 4t3 | Supercontig_12.7 | 3365564 | G | T | 0 | 1 | 38 | 26 | 0 | 26 | SNP | synonymous | GTG->GTT | V297V | | NCU02198 | > | hypothetical protein |
| Nc159 | 5t1 | Supercontig_12.4 | 978386 | C | G | 0 | 1 | 23 | 10 | 0 | 10 | SNP | intergenic | | | | | | |
| Nc159 | 5t1 | Supercontig_12.5 | 1607951 | T | C | 0 | 1 | 39 | 19 | 0 | 19 | SNP | UTR | | | | NCU04584 | > | hypothetical protein |
| Nc159 | 5t1 | Supercontig_12.6 | 910513 | G | GC | 0 | 0.9375 | 50 | 32 | 0 | 30 | INS | frameshift | | | | NCU04811 | < | hypothetical protein |
| Nc159 | 5t1 | Supercontig_12.7 | 1733870 | A | C | 0 | 1 | 53 | 28 | 0 | 28 | SNP | missense | CTG->CGG | L146R | gtp-24 | NCU06122 | < | guanine triphosphate binding-24 |
| Nc159 | 5t2 | Supercontig_12.1 | 1895511 | C | A | 0 | 1 | 41 | 31 | 0 | 31 | SNP | UTR | | | msp-13 | NCU01950 | > | pre-mRNA-splicing factor slu-7 |
| Nc159 | 5t2 | Supercontig_12.1 | 2238999 | G | T | 0 | 1 | 49 | 26 | 0 | 26 | SNP | missense | TGC->TTC | C630F | drh-20 | NCU01857 | > | DEAD/DEAH box RNA helicase |
| Nc159 | 5t2 | Supercontig_12.1 | 2812371 | C | A | 0 | 1 | 38 | 23 | 0 | 23 | SNP | missense | CAC->AAC | H861N | gla-2 | NCU02583 | > | Alpha-glucosidase |
| Nc159 | 5t2 | Supercontig_12.1 | 8296468 | C | A | 0 | 1 | 31 | 29 | 0 | 29 | SNP | missense | AGG->AGT | R123S | mox-3 | NCU03049 | < | monooxygenase-3 |
| Nc159 | 5t2 | Supercontig_12.2 | 2998109 | G | T | 0 | 1 | 44 | 47 | 0 | 47 | SNP | missense | GAC->GAA | D163E | rpn-6 | NCU01596 | < | 26S proteasome non-ATPase regulatory subunit 11 |
| Nc159 | 5t2 | Supercontig_12.3 | 3995452 | G | C | 0 | 1 | 52 | 24 | 0 | 24 | SNP | nonsense | TAC->TAG | Y235* | acon-2 | NCU00478 | < | aconidiate-2 |
| Nc159 | 5t2 | Supercontig_12.4 | 3514928 | C | G | 0 | 1 | 30 | 39 | 0 | 39 | SNP | missense | GTG->CTG | V200L | mdr-4 | NCU04391 | < | multi-drug resistance-4 |
| Nc159 | 5t2 | Supercontig_12.5 | 5815080 | C | A | 0 | 1 | 69 | 31 | 0 | 31 | SNP | intergenic | | | | | | |
| Nc159 | 5t2 | Supercontig_12.6 | 297902 | C | A | 0 | 1 | 39 | 19 | 0 | 19 | SNP | missense | GGG->TGG | G384W | apr-8 | NCU07063 | < | hypothetical protein |
| Nc159 | 5t2 | Supercontig_12.6 | 3647983 | G | T | 0 | 1 | 26 | 25 | 0 | 25 | SNP | intergenic | | | | | | |
| Nc159 | 6t1 | Supercontig_12.2 | 935003 | G | T | 0 | 1 | 39 | 42 | 0 | 42 | SNP | missense | GCC->TCC | A248S | fam-8 | NCU03372 | > | nonspecific lipid-transfer protein |
| Nc159 | 6t1 | Supercontig_12.2 | 4396932 | T | C | 0 | 1 | 46 | 42 | 0 | 42 | SNP | synonymous | CAA->CAG | Q176Q | cel-3 | NCU09473 | < | 3-ketoacyl-acyl carrier protein reductase |
| Nc159 | 6t1 | Supercontig_12.3 | 102394 | C | G | 0 | 1 | 49 | 31 | 0 | 31 | SNP | missense | GCA->GGA | A670G | | NCU09503 | > | hypothetical protein |
| Nc159 | 6t1 | Supercontig_12.4 | 528411 | C | A | 0 | 1 | 30 | 25 | 0 | 25 | SNP | synonymous | GCC->GCA | A959A | pks-3 | NCU04865 | > | polyketide synthase 3 |
| Nc159 | 6t1 | Supercontig_12.4 | 3948731 | C | G | 0 | 1 | 69 | 29 | 0 | 29 | SNP | missense | GGG->GCG | G1326A | | NCU08023 | < | cytohesin-2 |
| Nc159 | 6t1 | Supercontig_12.4 | 4967822 | G | C | 0 | 1 | 48 | 40 | 0 | 40 | SNP | missense | AAC->AAG | N994K | | NCU07029 | < | hypothetical protein |
| Nc159 | 6t1 | Supercontig_12.6 | 3348079 | GT | G | 0 | 0.846153846 | 42 | 26 | 0 | 22 | DEL | frameshift | | | | NCU05950 | > | hypothetical protein |
| Nc159 | 6t1 | Supercontig_12.6 | 3348081 | A | G | 0 | 0.956521739 | 41 | 23 | 0 | 22 | SNP | missense | AGG->GGG | R12G | | NCU05950 | > | hypothetical protein |
| Nc159 | 6t1 | Supercontig_12.6 | 3348084 | A | G | 0 | 1 | 41 | 23 | 0 | 23 | SNP | missense | AGG->GGG | R13G | | NCU05950 | > | hypothetical protein |
| Nc159 | 6t2 | Supercontig_12.1 | 3037951 | GC | G | 0 | 0.96969697 | 48 | 33 | 0 | 32 | DEL | intergenic | | | | | | |
| Nc159 | 6t2 | Supercontig_12.1 | 9026874 | T | G | 0 | 1 | 36 | 24 | 0 | 24 | SNP | missense | CTT->CGT | L294R | os-1 | NCU02815 | > | histidine kinase |
| Nc159 | 6t2 | Supercontig_12.2 | 1689233 | C | A | 0 | 1 | 36 | 16 | 0 | 16 | SNP | missense | CGT->CTT | R61L | | NCU05470 | < | hypothetical protein |
| Nc159 | 6t2 | Supercontig_12.2 | 2906482 | A | AC | 0 | 0.8125 | 12 | 16 | 0 | 13 | INS | intergenic | | | | | | |
| Nc159 | 6t2 | Supercontig_12.3 | 1378722 | G | C | 0 | 1 | 35 | 29 | 0 | 29 | SNP | intergenic | | | | | | |
| Nc159 | 6t2 | Supercontig_12.5 | 1450803 | C | A | 0.025641026 | 1 | 39 | 37 | 1 | 37 | SNP | missense | GGT->GTT | G137V | gh35-2 | NCU04623 | < | Beta-galactosidase |
| Nc159 | 6t2 | Supercontig_12.5 | 2921427 | A | G | 0 | 1 | 18 | 18 | 0 | 18 | SNP | intronic | | | oxa-1 | NCU01102 | > | mitochondrial export translocase Oxa1 |
| Nc159 | 6t2 | Supercontig_12.5 | 3766053 | C | A | 0 | 1 | 49 | 17 | 0 | 17 | SNP | missense | ACC->AAC | T81N | | NCU16999 | > | hypothetical protein |

| | | | | | | | | | | | | | | | | | | | |
|---|---|---|---|---|---|---|---|---|---|---|---|---|---|---|---|---|---|---|---|
| Nc159 | 6t2 | Supercontig_12.7 | 1634122 | G | T | 0 | 1 | 31 | 27 | 0 | 27 | SNP | UTR | | | drh-10/ | NCU06149/NCU17 | >/< | ATP-dependent RNA helicase dhh-1/hypothetical protein |
| Nc159 | 7t1 | Supercontig_12.1 | 474884 | G | A | 0 | 1 | 45 | 31 | 0 | 31 | SNP | intronic | | | | NCU07481 | > | morphogenesis protein |
| Nc159 | 7t1 | Supercontig_12.2 | 2425215 | T | C | 0.025641026 | 1 | 39 | 23 | 1 | 23 | SNP | intergenic | | | | | | |
| Nc159 | 7t1 | Supercontig_12.2 | 4206471 | C | T | 0 | 1 | 37 | 19 | 0 | 19 | SNP | intergenic | | | | | | |
| Nc159 | 7t1 | Supercontig_12.3 | 4563732 | C | A | 0 | 1 | 34 | 26 | 0 | 26 | SNP | intergenic | | | | | | |
| Nc159 | 7t1 | Supercontig_12.4 | 1487657 | A | C | 0 | 1 | 23 | 30 | 0 | 30 | SNP | missense | TAT->TCT | Y338S | | NCU06513 | > | hypothetical protein |
| Nc159 | 7t1 | Supercontig_12.4 | 2380931 | G | A | 0 | 1 | 30 | 24 | 0 | 24 | SNP | nonsense | CAG->TAG | Q557* | | NCU08331 | < | hypothetical protein |
| Nc159 | 7t1 | Supercontig_12.5 | 3373370 | A | C | 0 | 1 | 41 | 20 | 0 | 20 | SNP | intergenic | | | | | | |
| Nc159 | 7t2 | Supercontig_12.1 | 1415152 | G | GGA | 0 | 0.956521739 | 33 | 23 | 0 | 22 | INS | intergenic | | | | | | |
| Nc159 | 7t2 | Supercontig_12.1 | 7549393 | G | C | 0 | 1 | 40 | 18 | 0 | 18 | SNP | UTR | | | | NCU00687 | < | hypothetical protein |
| Nc159 | 7t2 | Supercontig_12.2 | 4047502 | C | A | 0 | 1 | 44 | 22 | 0 | 22 | SNP | intergenic | | | | | | |
| Nc159 | 7t2 | Supercontig_12.3 | 3994713 | GT | G | 0 | 0.967741935 | 41 | 31 | 0 | 30 | DEL | frameshift | | | acon-2 | NCU00478 | < | aconidiate-2 |
| Nc159 | 7t2 | Supercontig_12.6 | 859194 | G | T | 0 | 0.828571429 | 50 | 35 | 0 | 29 | SNP | intergenic | | | | | | |
| Nc159 | 7t2 | Supercontig_12.7 | 2876542 | A | G | 0 | 1 | 56 | 33 | 0 | 33 | SNP | intergenic | | | | | | |
| Nc159 | 8t1 | Supercontig_12.1 | 5491962 | A | G | 0 | 0.882352941 | 10 | 17 | 0 | 15 | SNP | intergenic | | | | | | *likely false-positive |
| Nc159 | 8t1 | Supercontig_12.3 | 4881015 | A | C | 0 | 1 | 39 | 22 | 0 | 22 | SNP | missense | ACC->CCC | T6P | ad-19 | NCU08568 | > | adenine-19 |
| Nc159 | 8t1 | Supercontig_12.3 | 5203062 | G | T | 0 | 1 | 37 | 19 | 0 | 19 | SNP | intergenic | | | | | | |
| Nc159 | 8t1 | Supercontig_12.4 | 977962 | T | G | 0 | 1 | 24 | 19 | 0 | 19 | SNP | intergenic | | | | | | |
| Nc159 | 8t1 | Supercontig_12.4 | 1053118 | G | T | 0 | 1 | 13 | 14 | 0 | 14 | SNP | intergenic | | | | | | |
| Nc159 | 8t1 | Supercontig_12.4 | 4931502 | C | A | 0 | 1 | 45 | 26 | 0 | 26 | SNP | UTR | | | vps17 | NCU07020 | < | vacuolar protein sorting-associated protein vps17 |
| Nc159 | 8t1 | Supercontig_12.5 | 6317475 | G | A | 0 | 1 | 60 | 28 | 0 | 28 | SNP | missense | ACC->ATC | T5I | | NCU07181 | < | hypothetical protein |
| Nc159 | 8t1 | Supercontig_12.6 | 3348056 | C | G | 0 | 1 | 35 | 31 | 0 | 31 | SNP | missense | AAC->AAG | N3K | | NCU05950 | > | hypothetical protein |
| Nc159 | 8t2 | Supercontig_12.1 | 3336083 | G | A | 0 | 1 | 36 | 32 | 0 | 32 | SNP | missense | CTT->TTT | L224F | | NCU02725 | < | hypothetical protein |
| Nc159 | 8t2 | Supercontig_12.1 | 4745500 | A | G | 0 | 1 | 58 | 32 | 0 | 32 | SNP | missense | AAG->AGG | K388R | | NCU03154 | > | Pantothenate kinase |
| Nc159 | 8t2 | Supercontig_12.1 | 6758806 | C | T | 0 | 1 | 53 | 28 | 0 | 28 | SNP | intergenic | | | | | | |
| Nc159 | 8t2 | Supercontig_12.1 | 7615373 | C | G | 0 | 1 | 48 | 33 | 0 | 33 | SNP | missense | TCG->TGG | S610W | | NCU00671 | > | beige/BEACH domain-containing protein |
| Nc159 | 8t2 | Supercontig_12.1 | 9026654 | G | A | 0 | 1 | 47 | 32 | 0 | 32 | SNP | missense | GGA->AGA | G519R | os-1 | NCU02815 | > | histidine kinase |
| Nc159 | 8t2 | Supercontig_12.5 | 1213066 | G | C | 0 | 1 | 23 | 10 | 0 | 10 | SNP | intergenic | | | | | | |
| Nc159 | 8t2 | Supercontig_12.5 | 1967191 | G | T | 0 | 1 | 62 | 24 | 0 | 24 | SNP | intergenic | | | | | | |
| Nc159 | 8t2 | Supercontig_12.5 | 2100422 | G | A | 0 | 1 | 52 | 24 | 0 | 24 | SNP | nonsense | CGA->TGA | R538* | | NCU03694 | < | hypothetical protein |
| Nc159 | 8t2 | Supercontig_12.6 | 2083100 | G | T | 0 | 1 | 36 | 25 | 0 | 25 | SNP | intergenic | | | | | | |
| Nc159 | 8t2 | Supercontig_12.7 | 2389132 | T | G | 0 | 1 | 18 | 10 | 0 | 10 | SNP | intergenic | | | | | | |

**Features found using custom script:**

| Ancestor | Evolved m | Scaffold | Bin # (*100 bp) | Total cov | Coverage of | Total coverage in | Coverage of alt | Type | Effect/Location | | Gene name | Gene code (NCU | Gene orientation | Comment |
|---|---|---|---|---|---|---|---|---|---|---|---|---|---|---|
| Nc159 | 8t2 | Supercontig_12.5 | 638 | 37.01 | 0.99 | 24.8 | 18.31 | INS | intergenic | 6bp insertion | | | | |
| Nc159 | 7t2 | Supercontig_12.1 | 14152 | 33.3 | 0 | 23.14 | 17.63 | | | | | | | |
| Nc159 | 7t2 | Supercontig_12.3 | 39947 | 41.93 | 0.37 | 34.69 | 23.96 | | | | | | | |
| Nc159 | 6t2 | Supercontig_12.1 | 30379 | 51.9 | 0 | 37.15 | 26.46 | | | | | | | |
| Nc159 | 6t1 | Supercontig_12.6 | 33481 | 45.33 | 2.06 | 25.83 | 20 | | | | | | | |
| Nc159 | 5t1 | Supercontig_12.6 | 9105 | 52.81 | 0.35 | 35.88 | 26.29 | | | | | | | |
| Nc159 | 4t1 | Supercontig_12.1 | 34376 | 48.75 | 0.91 | 19.31 | 15.02 | | | | | | | |
| Nc159 | 4t1 | Supercontig_12.7 | 42539 | 33.85 | 0 | 19.68 | 17.43 | DEL | intergenic | ~1370bp deletion | | | | |
| Nc159 | 1t2 | Supercontig_12.7 | 42543 | 16.74 | 0.25 | 18.02 | 16.75 | DEL | intergenic | ~980bp deletion | | | | |
| Nc152 | 9t2 | Supercontig_12.1 | 33366 | 58.06 | 1.75 | 32.92 | 20.29 | | | | | | | |
| Nc152 | 9t2 | Supercontig_12.2 | 37727 | 44.59 | 0.09 | 29.07 | 23.14 | | | | | | | |
| Nc152 | 16t3 | Supercontig_12.1 | 47322 | 60.07 | 0 | 26.74 | 15.07 | | | | | | | |
| Nc152 | 16t3 | Supercontig_10.21 | 58 | 148.52 | 5.52 | 115.12 | 69.79 | INS | intergenic | 31bp insertion | | | | |
| Nc152 | 16t3 | Supercontig_10.21 | 429 | 248.37 | 5.87 | 275.28 | 206.55 | | | | | | | |
| Nc152 | 16t2 | Supercontig_12.1 | 78740 | 54.21 | 0.72 | 24.29 | 18.02 | | | | | | | |
| Nc152 | 16t2 | Supercontig_10.21 | 58 | 148.52 | 5.52 | 133.37 | 90.69 | INS | intergenic | 31bp insertion | | | | |
| Nc152 | 16t2 | Supercontig_10.21 | 429 | 248.37 | 5.87 | 402.75 | 287.99 | | | | | | | |
| Nc152 | 16t1 | Supercontig_10.21 | 58 | 148.52 | 5.52 | 128.61 | 83.97 | INS | intergenic | 31bp insertion | | | | |
| Nc152 | 16t1 | Supercontig_10.21 | 429 | 248.37 | 5.87 | 324.7 | 239.61 | | | | | | | |
| Nc152 | 15t2 | Supercontig_12.1 | 68387 | 43.14 | 1.24 | 16.3 | 13.04 | DEL | frameshift | 41bp deletion | sin3-2 | NCU00897 | > | Sin3-like-2 |
| Nc152 | 15t2 | Supercontig_12.1 | 91120 | 42.5 | 0 | 34.4 | 20.55 | INS | frameshift | 1bp insertion | so | NCU02794 | > | Fso1 |
| Nc152 | 15t1 | Supercontig_12.4 | 15263 | 59.67 | 1.77 | 29.38 | 15.87 | INS | inframe | 6bp insertion | | NCU06525 | < | hypothetical protein |
| Nc152 | 15t1 | Supercontig_12.5 | 38626 | 41.29 | 0 | 20.06 | 12.71 | INS | intergenic | 6bp insertion | | | | |
| Nc152 | 14t2 | Supercontig_12.1 | 89268 | 70.55 | 0.9 | 15.81 | 13.41 | DEL | 2 partial and 8 c | ~39560bp deletior | NA/NA/NA/gh11-1/I | NCU02858/NCU0 | </</</</>/>/>/ | hypothetical protein/hypothetical protein/hypothetical protein/endo-1,4-beta-xylanase A/hypothetical protein/zinc finger transcription factor-35/Cytochrome P450/alcohol dehydrogenase-4/hypothetical protein/hypothetical protein |
| Nc152 | 14t2 | Supercontig_12.1 | 89334 | 51.09 | 0.28 | 18.93 | 15.21 | DEL | 1 complete gene | ~4680bp deletion | | NCU02843 | < | hypothetical protein |
| Nc152 | 14t2 | Supercontig_12.1 | 91120 | 42.5 | 0 | 18.06 | 12.16 | DEL | 2 partial gene de | ~22850bp deletior | so/NA | NCU02794/NCU0 | >/> | Fso1/Leashin Woronin body tether |
| Nc152 | 14t2 | Supercontig_10.21 | 110 | 160.84 | 2.3 | 283.38 | 146.52 | DUPL | intergenic | 234bp duplication | | | | *probable duplication |
| Nc152 | 14t1 | Supercontig_10.21 | 110 | 160.84 | 2.3 | 620.65 | 364.36 | DUPL | intergenic | 234bp duplication | | | | *probable duplication |
| Nc152 | 13t4 | Supercontig_12.1 | 91118 | 37.72 | 0 | 24.45 | 17.2 | DEL | 1 partial gene de | 937bp deletion | so | NCU02794 | > | Fso1 |
| Nc152 | 13t4 | Supercontig_12.6 | 3041 | 51.4 | 0 | 21.47 | 16.83 | DEL | 1 partial gene de | 76bp deletion | stk-49 | NCU07062 | < | protein kinase |
| Nc152 | 13t4 | Supercontig_12.7 | 27695 | 55.25 | 1.6 | 23.6 | 15.82 | | | | | | | |
| Nc152 | 13t3 | Supercontig_12.1 | 91119 | 41.37 | 0 | 21.61 | 17.89 | DEL | 1 partial gene de | 937bp deletion | so | NCU02794 | > | Fso1 |
| Nc152 | 13t3 | Supercontig_12.3 | 23195 | 43.35 | 1.78 | 30.21 | 22.16 | INS | inframe | 12bp insertion | | NCU00021 | < | hypothetical protein |
| Nc152 | 13t3 | Supercontig_12.6 | 3041 | 51.4 | 0 | 31.26 | 18.37 | DEL | 1 partial gene de | 76bp deletion | stk-49 | NCU07062 | < | protein kinase |
| Nc152 | 13t3 | Supercontig_12.7 | 27695 | 55.25 | 1.6 | 31.48 | 23.31 | | | | | | | |
| Nc152 | 13t2 | Supercontig_12.1 | 91119 | 41.37 | 0 | 21.83 | 17.67 | DEL | 1 partial gene de | 937bp deletion | so | NCU02794 | > | Fso1 |
| Nc152 | 13t2 | Supercontig_12.4 | 21260 | 54.58 | 0.67 | 30.94 | 19.08 | | | | | | | |
| Nc152 | 13t2 | Supercontig_12.7 | 27695 | 55.25 | 1.6 | 35.89 | 28.86 | | | | | | | |
| Nc152 | 12t2 | Supercontig_12.5 | 48529 | 50.67 | 0 | 31.44 | 22.94 | | | | | | | |
| Nc152 | 12t1 | Supercontig_12.5 | 25405 | 57.77 | 0.89 | 32.98 | 16.7 | INS | intergenic | 2bp insertion | | | | |
| Nc152 | 12t1 | Supercontig_10.21 | 429 | 248.37 | 5.87 | 553.86 | 406.91 | | | | | | | |
| Nc152 | 11t3 | Supercontig_12.4 | 24179 | 51.21 | 0 | 27.8 | 17.52 | | | | | | | |
| Nc152 | 11t3 | Supercontig_12.6 | 14786 | 56.72 | 8.26 | 28.62 | 23.52 | | | | | | | |
| Nc152 | 10t2 | Supercontig_12.3 | 39953 | 56.22 | 0.86 | 31.45 | 16.82 | DEL | 1 partial gene de | 64bp deletion | acon-2 | NCU00478 | < | aconidiate-2 |
| Nc152 | 10t1 | Supercontig_12.1 | 92144 | 50.07 | 3.35 | 30.66 | 24.06 | | | | | | | |
| Nc152 | 10t1 | Supercontig_12.2 | 9638 | 43.41 | 0 | 28.6 | 16.15 | | | | | | | |
| Nc152 | 10t1 | Supercontig_12.6 | 14786 | 56.72 | 8.26 | 25.06 | 19.5 | | | | | | | |

red colour indicates features already found by VarScan above
